# Evolving spike-protein *N*-glycosylation in SARS-CoV-2 variants

**DOI:** 10.1101/2023.05.08.539897

**Authors:** Sabyasachi Baboo, Jolene K. Diedrich, Jonathan L. Torres, Jeffrey Copps, Bhavya Singh, Patrick T. Garrett, Andrew B. Ward, James C. Paulson, John R. Yates

**Affiliations:** Department of Molecular Medicine, The Scripps Research Institute, La Jolla, California 92037, United States; Department of Integrative Structural and Computational Biology, The Scripps Research Institute, La Jolla, California 92037, United States; Department of Immunology and Microbiology, The Scripps Research Institute, La Jolla, California 92037, United States

**Keywords:** *N*-glycan, heterogeneity, SARS-CoV-2, COVID-19, spike-protein, variants, proteomics, mass spectrometry, vaccine, virus

## Abstract

Since >3 years, SARS-CoV-2 has plunged humans into a colossal pandemic. Henceforth, multiple waves of infection have swept through the human population, led by variants that were able to partially evade acquired immunity. The co-evolution of SARS-CoV-2 variants with human immunity provides an excellent opportunity to study the interaction between viral pathogens and their human hosts. The heavily *N*-glycosylated spike-protein of SARS-CoV-2 plays a pivotal role in initiating infection and is the target for host immune-response, both of which are impacted by host-installed *N*-glycans. Using highly-sensitive DeGlyPHER approach, we compared the *N*-glycan landscape on spikes of the SARS-CoV-2 Wuhan-Hu-1 strain to seven WHO-defined variants of concern/interest, using recombinantly expressed, soluble spike-protein trimers, sharing same stabilizing-mutations. We found that *N*-glycan processing is conserved at most sites. However, in multiple variants, processing of *N*-glycans from high mannose- to complex-type is reduced at sites N165, N343 and N616, implicated in spike-protein function.

## Introduction

The COVID-19 pandemic caused by SARS-CoV-2^1,2^ has highlighted the threat that zoonotic viruses pose to human health and socio-economic sustainability^3,4^. The current pandemic is one of the most severe humans have experienced, with a global death toll of 6.9 million and infection of more than 9.6% of global human population (https://covid19.who.int/, https://www.census.gov/popclock/). The original SARS-2 strain is believed to have begun as a moderately transmissive virus, infecting its first human host(s) in China in late 2019^5-7^. Since then, SARS-CoV-2 has infected the human population in surges, mutating frequently and evolving into increasingly more transmissive variants^4,8-13^ **(Figure 1A)**. Wuhan-Hu-1, the original SARS-CoV-2 strain, was the dominant strain throughout 2020, and has since produced 5 variants of concern (VoC), as defined by World Health Organization (WHO): Alpha, Beta, Gamma, Delta, and Omicron^14^. The Alpha variant was first detected in the United Kingdom, was highly transmissible, and quickly became the most dominant variant outside of South America and Africa from December 2020 until July 2021^15,16^. Almost simultaneously, Beta was detected in South Africa and became dominant in Africa^17^, and Gamma was detected in Brazil and became dominant in South America^18^. The highly virulent and transmissible Delta^19^ was first detected in India in mid-2021 and became the dominant variant until it was overtaken in January 2022 by the currently dominant Omicron^20^. GISAID (https://gisaid.org/) followed the spread of 2 WHO-recognized variants of interest (VoI), Mu^21,22^ (first detected in Colombia) and Lambda^23^ (first detected in Peru), which emerged in 2021 and partially overlapped with Gamma in South America.

**Figure 1:**
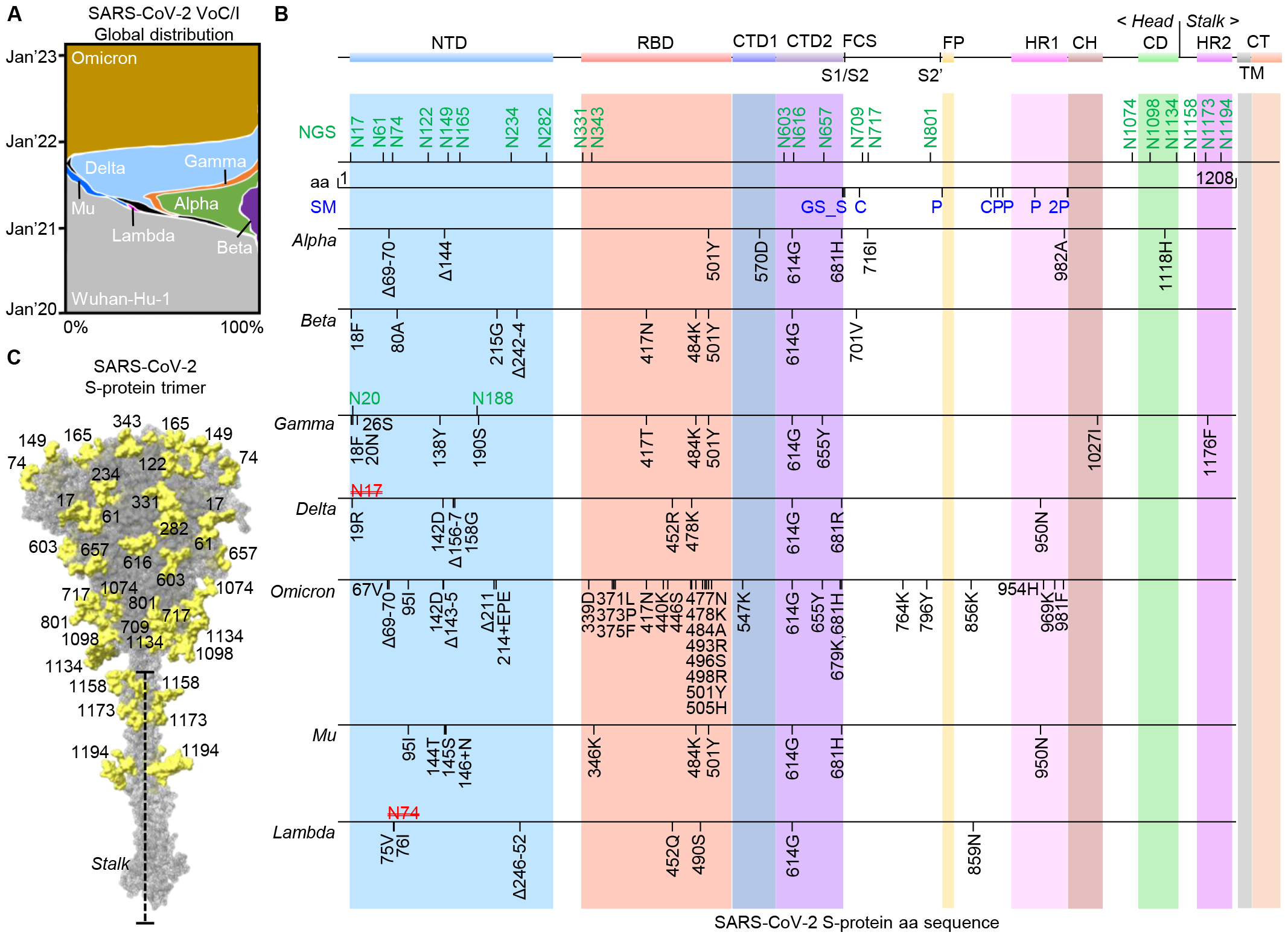
Mutational landscape of spike-protein from SARS-CoV-2 variants. (A) Global waves of SARS-CoV-2 variants over last 3 years and their temporal proportions (modified from nextstrain.org). (B) Critical domains along the amino acid (aa) length of spike-protein *viz*., N-terminal domain (NTD), Receptor-binding domain (RBD), C-terminal domains (CTD-1 and 2) of S1-protein, Fusion peptide (FP), Heptad repeats (HR-1 and 2), Central helix (CH), Connecting domain (CD), Transmembrane (TM) domain and Cytoplasmic tail (CT), with associated *N*-glycosylation sites (NGS, green), incorporated stabilizing mutations (SM, blue) and most frequent variant-specific mutations (deleted NGS in red). Furin cleavage site (FCS or S1/S2) and the subsequent cleavage site (S2’) for other cellular proteases is shown. (C) Structure of the full-length spike-protein trimer with representative *N*-glycans in yellow.

Thanks to decades of scientific research devoted to the improvement of vaccine technology and advances in an understanding of the immunology of viral infection (especially that of Coronaviruses, Human Immunodeficiency Virus (HIV) and Influenza)^24-33^ successful vaccines to counter SARS-CoV-2 infection were developed at an unprecedented pace^34-41^. These vaccines were designed against heavily glycosylated spike-protein (S-protein) trimer, commonly known as “spike”, anchored to the lipid-bilayer of the SARS-CoV-2 viral envelope^34,42^. This non-covalently associated trimeric form of spike-protein mediates virus entry by binding to its host cell-surface receptor, human ACE2 (Angiotensin converting enzyme 2)^43,44^. After fusing with a host cell, SARS-2 replicates its genetic material, and the host’s translation apparatus is hijacked to synthesize viral proteins^45,46^. The spike-protein is glycosylated with high mannose to complex type *N*-linked glycans at sequons encoded by Asn-X-Thr/Ser or NX(S|T), where X ≠ P^47^. *N*-glycans are added co-translationally by *en bloc* transfer from a lipid-containing intermediate to yield Glc_3_Man_9_GlcNAc_2_-Asn (Glc = Glucose, Man = Mannose, GlcNAc = *N*-acetylglucosamine). The monomeric spike proteins fold and assemble into a trimeric spike in the endoplasmic reticulum (ER) lumen^48-50^. Glycan addition at each NX(S/T) site occurs co-translationally through the *en bloc* transfer of a large high mannose oligosaccharide (Glc_3_Man_9_GlcNAc_2_). Trimming of outer Glc and Man residues is initiated in the ER, and further processing to complex type glycans occurs as the spike protein moves through the ER and Golgi complex (GC) enroute to the cell surface^48-50^. The degree of conversion of high mannose to complex type *N*-glycans at each glycosylation site is influenced by the dynamics of the underlying protein structure and the proximity of other *N*-glycans that can restrict access to the processing enzymes^51^. While most *N*-glycosylation sites on SARS-CoV-2 spike-protein, glycans are processed to complex type, there is minimal processing at a few sites (*e.g*., N234), resulting in high mannose type glycans in the mature spike protein.

Interest in site specific glycosylation of the spike-protein stems from the functional significance of these glycans in infection and the host immune response. The opening up of the spike-protein receptor binding domain (RBD) for initiating interaction with ACE2 on host cells, is shown to be regulated by glycans on N122, N165, N234 and N343^52-54^. In addition, *N*-glycans of viral spike glycoproteins shield the underlying protein structure facilitating escape from host immune responses^50,55,56^. Thus, for rational design of vaccines capable of offering protection against evolving SARS-CoV-2 variants, it is important to assess the *N*-glycan landscape on the SARS-CoV-2 spike^57-61^. Multiple critical mutations that have appeared on spike-protein of SARS-CoV-2 variants, are known to facilitate escape from humoral immunity^62,63^, and some of these mutations might be responsible for a change in the *N*-glycan landscape of the spike, partially explaining the immune escape strategies of SARS-CoV-2 variants^9,42^.

Here, we present a systematic evaluation of the *N*-glycan landscape of the spike-protein from the original SARS-CoV-2 strain and 7 of its WHO-defined major variants (Alpha, Beta, Gamma, Delta, Omicron, Mu, and Lambda). Although a number of studies, using mass spectrometry (MS)-based glycoproteomic approaches, have compared glycosylation of SARS-COV-2 variant spike-proteins^64-69^, with the glycosylation of the original Wuhan-Hu-1 spike-protein^51,57,60,70-76^, they have examined a smaller subset of variants and there is substantial differences with regards to the degree of glycan processing and glycan occupancy at each glycosylation site **(Figure S1)**. Such variation can be expected for sites that have insufficient glycopeptide sampling to establish an accurate representation of the “*N*-glycan demographic”.

Here we have compared multiple SARS-CoV-2 variant spike-proteins, to make critical inferences from site-specific *N*-glycosylation changes with high statistical confidence – a property unique to our DeGlyPHER approach, owing to hugely enhanced peptide sampling due to Proteinase K digestion along with Endo H/PNGase F sequential deglycosylation^49,77-79^. We assess the impact on glycosylation of mutations needed to produce high-yielding and stable “soluble spike-protein trimers” for vaccines. We compare our results with the *N*-glycan landscape of the recombinant spike manufactured by Novavax (NVX-CoV2373), used as a vaccine against SARS-CoV-2^59^, which has been recently authorized by United States Food and Drug Administration (US FDA) for emergency use.

## Results

### Analysis of *N*-glycan landscape on spike-protein

The spike-protein of Wuhan-Hu-1 is 1,273 amino acids long and *N*-glycosylated (we exclusively analyze *N*-glycans in this study) at 22 potential *N*-glycosylation sites (NGS)^57^ **(Figure 1B)**. The spike trimer is comprised of 3 protomers and has an apical broad head consisting of the receptor binding domain (RBD) surrounded by the N-terminal domain (NTD) at its base, and tapers down to a long stalk^34,53,80^ **(Figure 1C)**. The spike head of a single spike-protein protomer is >1,100 amino acid long, populated by 8 NGS in NTD, 2 NGS in RBD and another 9 NGS. The 10 NGS in NTD and RBD are some of the most extensively studied glycosylation sites^52,53,81,82^. The spike stalk consists of <150 amino acids per protomer, populated by only 3 NGS.

We used recombinantly expressed SARS-CoV-2 spike (M1-Q1208) in cultured human cells. These proteins lacked the transmembrane (TM) and C-terminal (CT) domains but included C-terminal tags to aid in trimerization and affinity purification. The Furin cleavage site (FCS, S1/S2) was abrogated (682RRAR substituted with 682GSAS) to stabilize the spike **(Figure 1B, Supplementary Tables 1,2)**^83^. The structural integrity and purity of the spikes in their prefusion state was validated with size-exclusion chromatography (SEC) and negative-stain electron microscopy (NS-EM) **(Figure S2)**. We employed the DeGlyPHER^77,79^ approach that uses sequential deglycosylation followed by bottom-up MS-based proteomics to resolve the site-specific *N*-glycan heterogeneity on spikes from 7 SARS-CoV-2 variants and the Novavax full-length spike-protein (NVX-CoV2373; expressed in Sf9 cells *i.e*., insect cells) and compared them with the Wuhan-Hu-1. DeGlyPHER allowed us to broadly resolve the abundance of high-mannose (including hybrid) and complex glycans at each NGS and to estimate glycan unoccupancy^79^.

### Original SARS-CoV-2 strain/Wuhan-Hu-1

Wuhan-Hu-1 is believed to have jumped from an undetermined animal reservoir(s) to humans at the end of 2019^5,6^, and its spike is one of the of the most studied protein structures in recent times^80^. Most of the 22 NGS on spike-protein are predominantly complex type; only N122, N234, N603, N709, N717, N801 and N1074 show a majority of high mannose-type glycans **(Figure 2A)**. The results provide a robust dataset for detailed comparison among 7 spike proteins of SARS-CoV-2 VoC/Is defined by WHO. The site-specific glycan distribution observed for the Wuhan-Hu-1 spike protein is largely consistent with several reports from the Crispin group using a traditional glycoproteomic approach^57,64,76^. More significant differences have been reported by Shajahan *et. al*.^65,72^, likely the result of a workflow that provides insufficient sampling at each site to quantitate relative proportions of the major glycan species **(Figure S1)**. We analyzed 3 different versions of the spike-protein that were mutated to stabilize the immunogen in its prefusion state **(Figure 1B)**. The first version, 2P, contains 2 residues (K986 and V987) that were substituted to proline and this mutant is used for most of the currently authorized SARS-CoV-2 vaccines^29,59^. The second version, 6P (or HexaPro), has 4 additional proline substitutions at positions F817, A892, A899, and A942 which help with stability and increase protein yield^83^. The third version, 6P-Mut7, adds an additional interprotomer disulfide bridge at positions V705C/T883C, thus further stabilizing the immunogen while keeping the RBD flexible^84^. When we compared the glycan distribution pattern between 2P, 6P, and 6P-Mut7, the 6P/6P-Mut7 mutants exhibited a significantly higher proportion of high mannose glycans at N603, N801 and N1074 relative to the 2P mutant, but there were no statistically significant differences between 6P and 6P-Mut7 mutants, **(Figures 2B,C,D)**. We see that different stabilizing mutations can differentially impact the glycosylation pattern on spikes, which could complicate comparison of variant spikes with differing stabilizing mutations. With this in mind, we only made glycosylation comparisons between spikes that were produced using the same stabilizing mutations (6P-Mut7 when comparing all variants in this study with Wuhan-Hu-1; 6P only when comparing Delta and Omicron with Wuhan-Hu-1; 2P when comparing Wuhan-Hu-1 and NVX-CoV2373).

**Figure 2:**
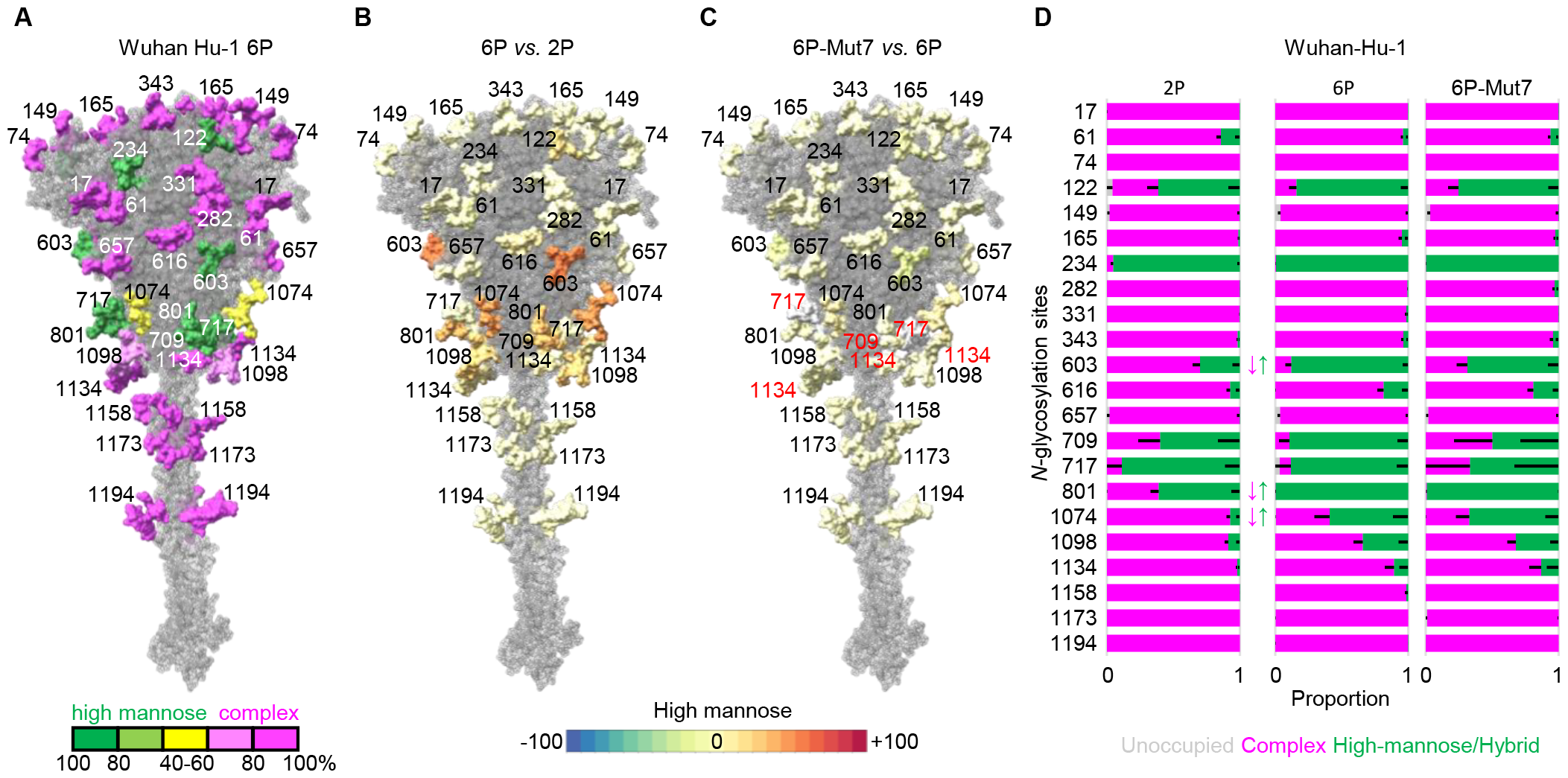
Comparing glycosylation changes on Wuhan-Hu-1 spike-protein owing to stabilizing mutations. (A) *N*-glycan heterogeneity determined on 6P version with dominant proportion color-coded from high mannose (green) to complex (magenta). A percentage-point scale signifying increase (towards red) or decrease (towards blue) in high mannose *N*-glycans, when performing site-specific comparison between 6P and 2P (B), showing significantly reduced *N*-glycan processing at N603, N801 and N1074; and 6P-Mut7 and 6P (C), showing no significant site-specific changes (Poor sampling of peptides mapping N709, N717 and N1134 in 6P-Mut7, numbered in red). (D) Bar-graphs demonstrating site-specific differences in glycosylation, if any, between 3 differently-stabilized mutants. *N*-glycosylation states are color-coded. Error bars represent mean–SEM. Any significant difference (BH-corrected p-value <0.05) in proportion of a certain *N*-glycosylation state at an NGS is represented by color-coded ↕.

### SARS-CoV-2 variants of concern/interest

For the first year of the COVID-19 pandemic the only mutation acquired by the SARS-CoV-2 spike-protein that could be positively selected to establish stable VoC/I was D614G^9,85^. The N501Y mutation in the spike RBD established 3 new VoCs, Alpha (501Y.V1 or B.1.1.7 or 20I), Beta (501Y.V2 or B.1.351 or 20H) and Gamma (501Y.V3 or P.1 or 20J)^86^. N501Y, which is reported to enhance binding affinity of spike RBD to human ACE2, is shared by Alpha, Beta and Gamma^87^. The other spike RBD mutations include E484K on both Beta and Gamma, and K417N on Beta and K417T on Gamma. The spike NTD mutations on Alpha (del69-70 and del144), Beta (L18F, D80A, D215G and del242-244) and Gamma (L18F, T20N, P26S, D138Y and R190S) are also known to affect neutralization^88^. None of the mutations in the Alpha or Beta resulted in gain or loss of an NGS, but there were notable changes in glycan processing at several NGS. In the Alpha spike-protein, there were significantly more complex glycans at N122, and more high mannose glycans at N616 **(Figure 3A)**. In the Beta spike, there were more high mannose glycans at N165 and N616 **(Figure 3A)**. The biggest changes in glycosylation occur in the Gamma spike-protein. Two of mutations resulted in a gain of NGS, T20N creating N20, and R190S creating N188. In the Gamma spike-protein **(Figure 3A)**, the new NGS at N20 contained predominantly complex type glycans, and the adjacent N17 NGS which was largely unoccupied. Moreover, the newly created NGS at N188 is close to N17 and N20 and had largely high mannose glycans. In contrast, glycan composition at N603 and N801 shifted significantly to complex type glycans.

**Figure 3:**
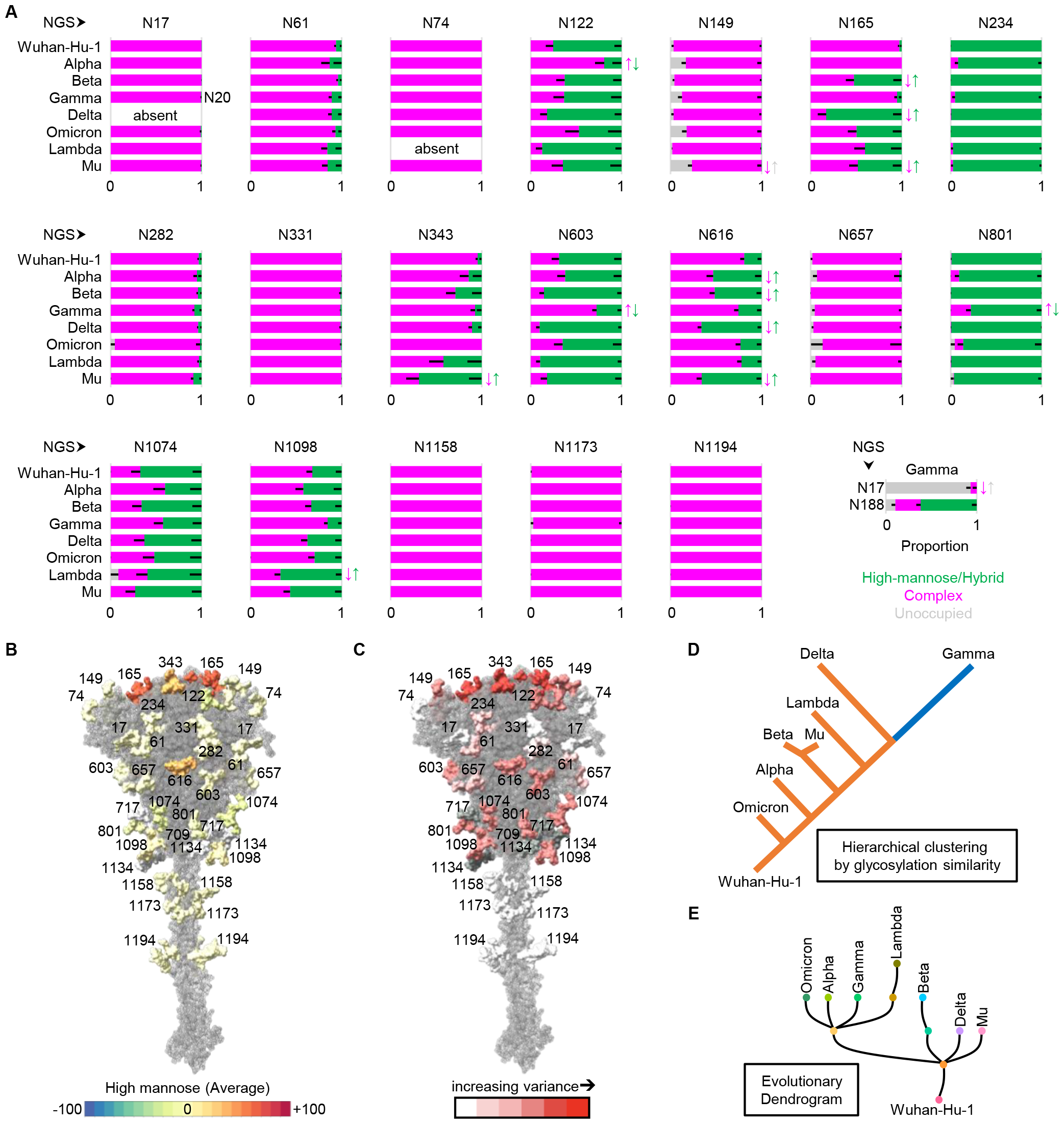
Site-specific comparison of SARS-CoV-2 6P-Mut7 spike-protein trimer from VoC/I with Wuhan-Hu-1. (A) Bar-graphs demonstrating site-specific differences in glycosylation, if any. N17 (glycosylation pattern transferred to N20 in Gamma), N61, N74, N149, N282, N331, N657, N1158, N1173 and N1194 are consistently occupied with mostly complex *N*-glycans across all variants (N17 and N74 absent in Delta and Lambda, respectively). N188 (only in Gamma), N234 and N801 are consistently occupied with mostly high mannose *N*-glycans across all variants. Across all variants, N1074 and N1098 are occupied by a more “even” distribution of high mannose and complex *N*-glycans. Variant specific changes in glycosylation pattern are most apparent at N122, N165, N343, N603 and N616. Across variants, NGS are mostly occupied with *N*-glycans. *N*-glycosylation states are color-coded. Error bars represent mean–SEM. Any significant difference (BH-corrected p-value <0.05) in proportion of a certain *N*-glycosylation state at an NGS, compared to the corresponding state in Wuhan-Hu-1, is represented by color-coded ↕. (B) A percentage-point scale signifying increase (towards red) or decrease (towards blue) in high mannose *N*-glycans, averaging across all variants, at each NGS, demonstrating highest shifts at N165, N343 and N616. (C) Variance in glycosylation state across all variants and Wuhan-Hu-1, with highest variance seen at N165 and N343. (D) Hierarchical clustering of SARS-CoV-2 variants and Wuhan-Hu-1 by similarity in glycosylation state shows Omicron is most similar to Wuhan-Hu-1, and Gamma and Delta the most dissimilar. (E) An evolutionary dendrogram of SARS-CoV-2 variants based on GISAID data considering nucleic acid mutations (covariants.org). None of the above analyses consider N709, N717 and N1134 owing to inconsistent sampling.

The Delta VoC (B.1.617.2 or 21A) emerged in 2021, spreading globally and replacing the predominance of 501Y lineage^19^. D614G is retained in the Delta spike-protein as well. In the Delta spike-protein, T19R mutation in NTD caused the loss of N17 NGS. There was a significant shift to high mannose glycans at N165 and N616 in both 6P and 6P-Mut7, and additionally at N343 in only 6P version. **(Figures 3A;4A,C**).

**Figure 4:**
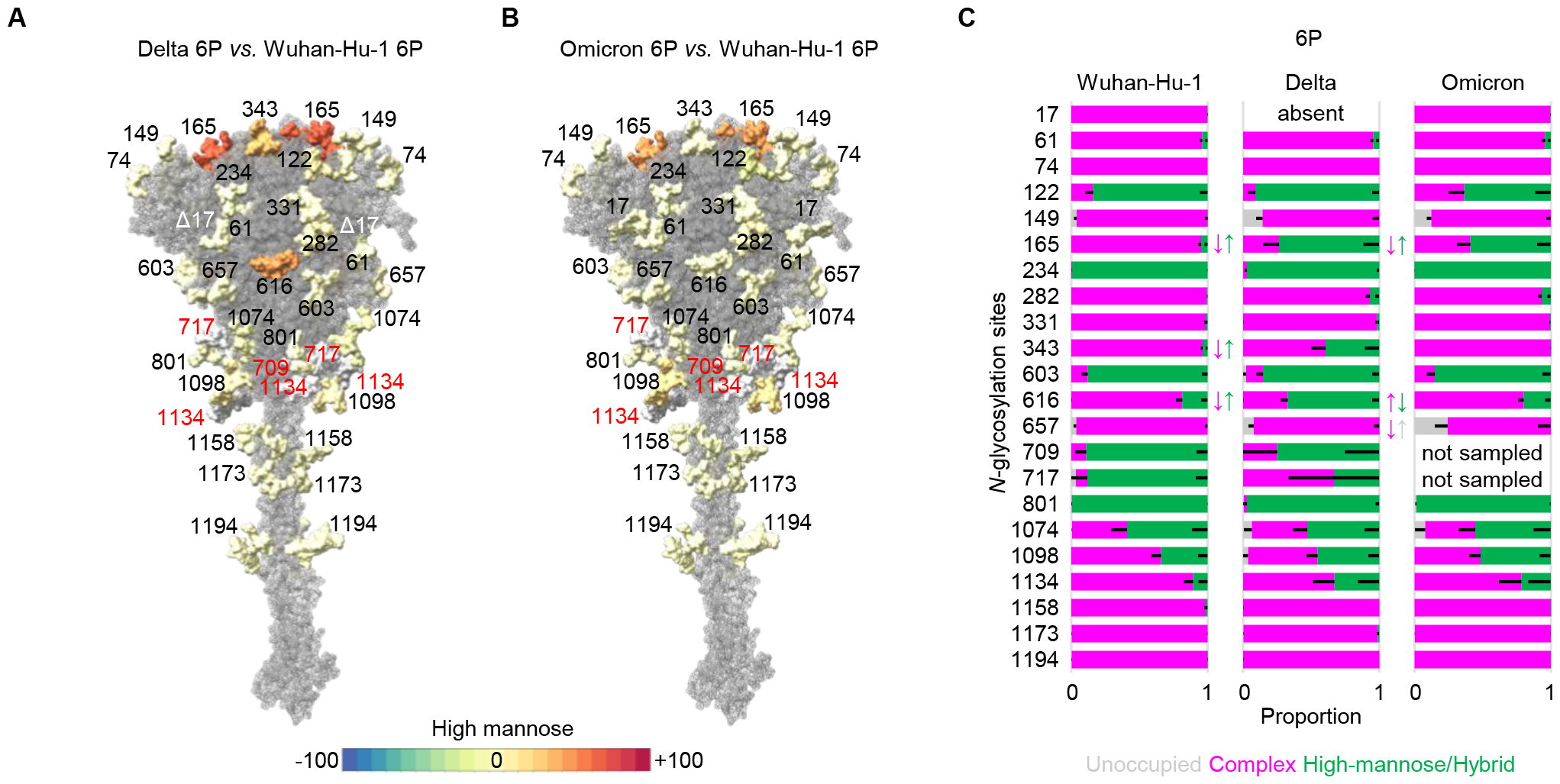
Site-specific comparison of SARS-CoV-2 6P spike-protein trimer from Wuhan-Hu-1, Delta and Omicron. A percentage-point scale signifying increase (towards red) or decrease (towards blue) in high mannose *N*-glycans, when performing site-specific comparison between Delta and Wuhan-Hu-1 (A), showing significantly reduced *N*-glycan processing at N165, N343 and N616; and Omicron and Wuhan-Hu-1 (B), showing prominently reduced *N*-glycan processing at N165, but not statistically significant. (C) Bar-graphs demonstrating site-specific differences in glycosylation, if any, between Wuhan-Hu-1 and Delta, and Delta and Omicron. *N*-glycosylation states are color-coded. Error bars represent mean–SEM. Any significant difference (BH-corrected p-value <0.05) in proportion of a certain *N*-glycosylation state at an NGS is represented by color-coded ↕. The significant shift from complex *N*-glycans in Wuhan-Hu-1 to high mannose *N*-glycans in Delta at N165, N343 and N616, reverts from Delta to Omicron.

The Omicron VoC (B.1.1.529 or 21M) replaced the Delta and spread globally in 2022, and Omicron subvariants still dominate^20^. The Omicron spike-protein hosts at least 33 mutations, none of which caused gain or loss of any NGS **(Figure 1B)** (though the newer Omicron subvariants are missing N17^89^). D614G is retained in the Omicron spike-protein. Despite this large number of mutations, when we compared the Omicron variant spike to spike from Wuhan-Hu-1 **(Figures 3A;4B,C)**, there was no significant shift in site-specific glycan processing. However, we note a shift from high mannose to complex glycans at N122 and *vice-versa* at N165. Additionally, we observed higher unoccupancy at N149 and N657.

The spread of Mu (B.1.1.621 or 21H) and Lambda (C.37 or 21G) VoIs was limited to South America^21,22^. The Mu spike-protein hosts 10 mutations while Lambda spike-protein hosts 7 mutations, the D614G mutation is retained in both **(Figure 1B)**, and include some of the known mutations (at L452, E484, N501 and P681) that help SARS-CoV-2 escape neutralization^21,22^. In the Mu spike-protein, there was a significant shift to high mannose glycans at N165, N343 and N616, while glycan unoccupancy increased significantly at N149 **(Figure 3A)**. In the Lambda spike-protein, T76I mutation in NTD caused the loss of N74 NGS and there was a significant shift to high mannose glycans only at N1098, though a similar trend was observed at N165 and N343 also **(Figure 3A)**.

### S2-protein

The post-fusion S2-protein structure that extends from the CT at its base to fusion peptide (FP) (or beyond if S2’ cleavage has not occurred) at the top^75,90^, hosts 3 stalk NGS (N1158, N1173 and N1194) in addition to the 5 NGS (N709, N717, N801, N1074, N1098 and N1134) that are structurally conserved in SARS-CoV^91^. The 5 NGS from N1098 to N1194 are spaced at almost regular intervals of 40Å^75,90^, suggesting that glycans at these NGS and at N1074 (assuming S2’ cleavage) are involved in “caging” the degrees of freedom of the stalk when the viral and host membranes are undergoing fusion^92^. We observed a statistically significant increase in high mannose glycans at N1098 in Lambda **(Figure 3A)**. Glycans anchored to N1098 on the apical rim of the post-fusion S2-protein^75,90^ could be important determinants of SARS-CoV-2’s infectivity, as observed in bronchial cells expressing ACE2 when N1098 is occupied with high-mannose *N*-glycans^93^. In this post-fusion S2-protein structure, N1074, N709 and N717 sit just below the apical rim as it extends into the Connecting domain (CD) that hosts N1134^75,90^.

The stalk of the spike is highly conserved across Beta-coronaviruses^94^, and thus it provides a promising target for immune intervention^95^. The 3 NGS (N1158, N1173 and N1194) on the stalk of the SARS-CoV-2 spike are fully occupied by complex glycans in all the variants **(Figure 3A)**. Complex glycans on these 3 NGS are known to massively shield the stalk at the flexible hinges that are prominent epitopes for immune recognition by antibodies^53,96^. Their conservation across evolving SARS-CoV-2 variants suggests that there is no apparent selection pressure from host immunity to modify the glycan shield around the stalk.

### NTD

NTD, which forms a whorl around the RBD, has 8 NGS (N17, N61, N74, N122, N149, N165, N234 and N282), the most of any of the structural domains in SARS-CoV-2 spike. Five of these NGS (N17, N61, N74, N234 and N282) have highly conserved glycan distributions across variants **(Figure 3A)**. Glycans at N17 are difficult to resolve on the spike structure using cryo-EM since it is in the N-terminal unstructured region^96^, except when stabilized by antibodies^80,81^. N17 is fully occupied by complex glycans **(Figure 3A)**, and this trend is conserved across all variants except Gamma, where it is mostly unoccupied, and in Delta and newer subvariants of Omicron, where it is absent^89^. In Gamma, the newly created N20 is known to be glycosylated and exterior to N17 on the N1 loop^81,97-99^, thus explaining why we observe it to be fully occupied by complex glycans, functionally replacing the adjacent N17, possibly impacting NTD antigenicity^68^. N17, along with N74, N122 and N149 border a multi-epitope “neutralization supersite” on NTD^81,100^. Absence of N17 in Delta is known to reduce neutralization by antibodies^97,100,101^ and absence of N74 (on N2 loop) in Lambda is known to increase infectivity (the spike-protein we used for Lambda did not include the new N254 NGS in N5 loop which is associated with reduced neutralization by antibodies^102-104^). N74 is fully occupied by complex glycans across all variants except Lambda, where it is absent **(Figure 3A)**. N122 is mostly occupied by high mannose glycans across variants, except in Alpha where N122 is significantly more occupied by complex glycans **(Figure 3A)**. N122 and N165 are proposed to affect an “RBD-up” conformation^54^. When the spike is in an “RBD-up” conformation^34^, glycans on N122 are predicted to significantly clash with the highly neutralizing S309 monoclonal antibody binding to its epitope in the “open” RBD. However, its binding is not expected to be affected^105,106^; indeed, no significant loss in neutralization of Alpha by S309 has been reported^107,108^, even though we see a shift in the type of glycans on N122 compared to Wuhan-Hu-1 **(Figure 3A)**. We found N149 (on surface-accessible N3 loop) to be mostly occupied by complex glycans **(Figure 3A)**. When these glycans include GlcNAc_2_Man_3_, N149 is reported to interact with host cell-surface lectin receptors, enhancing ACE2/spike-protein interaction in lung tissue and consequently driving cell-fusion and syncytia formation that is known to facilitate infection^109,110^. We observed that N149 is almost completely occupied by glycans in variants where the N3 loop is not mutated (Beta and Lambda), while the glycan unoccupancy at N149 is most significant on Mu **(Figure 3A)**. The glycosylation pattern on N165 shifts from almost completely complex type in Wuhan-Hu-1 virus to high mannose glycans in all variants except Alpha and Gamma **(Figure 3A)**. The inferred steric occlusion of N165 in most variants (the shift being highest in Delta) is highly relevant given the role of glycans at N165 and N234 in modulating conformational transitions of RBD and its stabilization in an “up” conformation and the RBM (receptor binding motif) is shielded in a “down” conformation^53,111^. The near-complete occupancy at N234 with high mannose glycans and at N282 with complex glycans is conserved across all variants we studied **(Figure 3A)**, with N282 strategically positioned in close proximity to the FP^82^ and also implicated in formation of dimeric spike-trimers^59^. In the Gamma variant only, N188 is at the C-terminal edge of the N4 loop, and we find it to be mostly occupied with high mannose glycans **(Figure 3A)**, which are reported to increase the glycan shielding effect^64^. Emerging mutations in recent Omicron subvariants, such as H245N mutation in BA.2.86 lineage, causes gain of N245 as NGS, and occurs along with loss of N17 (https://gisaid.org/hcov19-mutation-dashboard/), both in close proximity on NTD, possibly suggesting a functional role of glycans at this newly-emerged NGS.

### RBD

RBD, which physically interacts with ACE2 and initiates infection, hosts only 2 NGS (N331 and N343). Even in the heavily mutated RBD of Omicron (15 mutations), N331 and N343 are conserved and no new NGS arise. However, some emerging mutations in recent Omicron subvariants affect known *N*-glycosylation sites and/or create new ones (https://gisaid.org/hcov19-mutation-dashboard/) on spike-protein^112^. K529N mutation in an Omicron XBB.1.5 subvariant spike-protein RBD creates a new glycosylation site at the RBD-hinge, an otherwise conserved region^113^, close to N331, and its glycosylation can potentially regulate RBD “up/down” dynamics and indirectly reduce access to RBD epitopes^114,115^. N331 is populated almost completely with complex glycans across all variants we studied **(Figure 3A)**. N331 and N343 glycans are reported to shield the RBM in a “down” conformation^53^. We observed that N343 is occupied almost completely with complex glycans in the Wuhan-Hu-1 strain, as well as in some of the variants, but a shift towards high mannose glycans was seen in Lambda, Beta, and this shift was statistically significant in Mu and Delta (for 6P versions) variants **(Figures 3A;4A,C)**. These changes are highly relevant because glycans at N343 are implicated in pushing the RBD “up” into an “open” conformation^52^, and is targeted by neutralizing antibodies^106,108^.

### S1-protein CTD

The NTD-associated subdomain-2 of S1-protein *i.e*., CTD-1 and CTD-2 **(Figure 1B)** hosts 3 NGS (N603, N616 and N657). These 3 NGS, along with N61 and N801 in the S2-protein, are proximal to the proteolysis sites S1/S2 and S2’^116^. The glycans anchored to these NGS may affect access to Furin-like proteases through glycan holes around S1/S2, S2’ and FP^117^. Though we abrogated the S1/S2 cleavage site in the spike-proteins used in this study, any fluctuation in glycosylation state at these NGS could provide evidence for an interaction of the S1/S2 region with glycans on N61/N603^118^, and glycans on all 5 NGS that affect spike incorporation on virions and their capability to infect cells^116^. We observed a significantly higher proportion of high-mannose glycans on N603 and N801 in the 6P and 6P-Mut7 versus the 2P versions of Wuhan-Hu-1, as well as in in all variants we studied except Gamma, where complex glycans are significantly higher at N603/N801 **(Figure 3A)**. We find N61 and N657 to be almost completely occupied with complex glycans across all variants we studied **(Figure 3A)**.

N616 is adjacent to D614G, which is conserved in all SARS-CoV-2 variants that have emerged since Wuhan-Hu-1. The D614G mutation is known to have significantly increased SARS-CoV-2 infectivity by reducing S1-shedding, thus ensuring that the spike is intact and available for receptor binding^119-121^. The stabilization of spike in D614G variants is attributed to the loss of an interaction between G614 in one protomer and a region immediately downstream of FP (amino acids 823-862) in a neighboring protomer, and the effect this plays on an adjacent loop (loop 630) that is proposed to play a critical role in modulating/regulating spike structure for viral fusion with host^59,90,122,123^. Compared to Wuhan-Hu-1 where N616 is mostly occupied with complex glycans, Alpha, Beta, Delta, and Mu show significant shifts towards high-mannose glycans **(Figure 3A)**. The absence of this shift maybe explained by the FP-proximal T859N mutation in Lambda and N856K mutation in Omicron, which could possibly counter the loss of inter-protomer interaction caused due to D614G^124-127^. The change in glycosylation state at N616 in variants can contribute to or be affected by the structural dynamics of D614G^70^. This relationship has been postulated^85,128^, but not systematically evaluated.

## Discussion

The COVID-19 pandemic is driven by evolving variants of SARS-CoV-2, which are characterized by critical mutations that cause changes in the structure of the spike, hence affecting infectivity^129^. These structural modifications cause and are also affected by *N*-glycosylation changes. Using the 6P-Mut7 form of spike-protein from 7 WHO-defined SARS-CoV-2 variants, we present here a comprehensive analysis of changes in the site-specific *N*-glycan processing relative to Wuhan-Hu-1 using our highly-sensitive DeGlyPHER approach^77,79^ **(Figures 3A,B,C)**. We benchmarked the *N*-glycosylation pattern on Wuhan-Hu-1 6P-Mut7 spike-protein to its 2P and 6P versions that have been previously studied^57,59,60,65,72^. We note that the *N*-glycosylation pattern remains similar for most sites across these 3 mutant versions. However, there are a few site-specific differences (*e.g*., at 603, 1074, 1098) **(Figure 2)**, so we have taken care to compare SARS-CoV-2 variant spike-proteins using the same stabilizing mutations to ensure that any additional differences are due to variant mutations.

Using stabilized soluble spike-protein trimers hosting known variant-specific mutations, we observed that many NGS in NTD, RBD and stalk (N17/N20, N61, N74, N282, N331, N1158, N1173, N1194) exhibited near-complete occupancy with complex glycans, while N234 is occupied with high mannose glycans **(Figure 3A)**. Glycosylation changes were prominent **(Figure 3A)** at many NGS shown/predicted to be directly involved in: [1] RBD dynamics (N122, N149, N165 and N343)^53,54,81,109-111^, [2] proteolysis/spike-stabilization by virtue of their proximity to S1/S2 and/or S2’ (N603, N616 and N801)^70,85,116,118^, and [3] stabilization of post-fusion functions of S2-protein (N1098)^75,90,93^. Newby *et. al*.^64^ have made similar observations using 6P-stabilized spikes from 4 VoCs (Beta, Gamma, Delta and Omicron) using traditional MS-based glycoproteomic approaches. However, the results of Newby *et. al*. differ at a number of sites from those reported by Shajahan *et. al*.^65^ for 5 VoCs (Alpha, Beta, Gamma, Delta and Omicron) and Wang *et. al*.^66^ for 4 VoCs (Alpha, Beta, Gamma and Delta) using similar MS-based traditional glycoproteomic approaches. Moreover, in these previous studies, the results for the glycosylation pattern of the benchmark Wuhan-Hu-1 spike protein, against which the glycosylation pattern of variant spikes are being compared, also differ at multiple sites^57,64-66,70,72,76^ **(Figure S1)**. For example, using the same 6P-stabilized version of the Wuhan-Hu-1 spike one study found complete glycan occupancy at site N1173^76^ while another found no glycan occupancy^64^. Such variation is most likely explained by insufficient glycopeptide sampling per site in some studies, which we believe points to a strength of DeGlyPHER for more confident site-specific comparative analyses from extensive sampling at each site.

While our results on changes in glycosylation of variants largely agree with those of Newby *et. al*.^64^, there are several notable differences. While we concluded that several variants showed significantly increased high mannose at N165 relative to Wuhan-Hu-1, Newby *et. al*. ^64^ concluded there was little change as a result of greater high mannose in their analysis of Wuhan-Hu-1 and vice-versa for N61 where we do not observe any significant change in glycosylation **(Figures 3A;S1)**. Also, glycan unoccupancy reported by Newby *et. al*.^64^ was substantial at N331 (in Beta), N657 (in Wuhan-Hu-1, Beta, Gamma, Delta and Omicron), N1074 (Gamma), N1158 (Gamma), N1173 (in Wuhan-Hu-1, Beta, Gamma, Delta and Omicron; almost never occupied by glycans) and 1194 (in Beta, Gamma, Delta and Omicron), while we find all these sites to be almost completely occupied with glycans across all variants analyzed here **(Figures 3A;S1)**. Sparse peptide sampling at N709, N717 and N1134 **(Figure S3A)** precludes us from reading too much into the observed glycosylation pattern at these sites **(Figure S3A)**, though in all variants we studied, glycans decorating N1134 are mostly complex type **(Figure S3B)** and glycans at N709 and N717 on Wuhan-Hu-1 spike (2P/6P stabilized mutants, where we recorded sufficient peptide sampling) are mostly high mannose type **(Figures 3A;S1)**, as also reported by Newby *et. al*.^64^. The sparse sampling that we recorded may be attributable to structural constraints that limit protease access since we avoid harsh denaturants in our sample preparation. Alternatively, peptides mapping to these regions might have reduced ionization in the mass spectrometer^77,79^. Speculating from structural analysis of post-fusion spike^90,130^, glycans on N709/N717 may possibly affect proteolysis/spike-stabilization, and glycans on N1134 may play a role in post-fusion S2-protein stability.

Using a hierarchical clustering analysis for similarity in glycosylation pattern among shared and “well-sampled” NGS in 6P-Mut7, we see that despite the huge number mutations in Omicron, it is glycosylated more like Wuhan-Hu-1 than other variants, with Gamma and Delta being the most different **(Figure 3D)**. This hierarchical cluster differs from the inferred evolutionary pattern of the SARS-CoV-2 variants determined by nucleic acid mutation analysis (**Figure 3E**, modified from https://covariants.org/), suggesting that selective pressure on glycan changes is more dependent on higher-order structure than primary sequence of a protein, as evident from Wuhan-Hu-1 and Omicron spike-protein trimers being structurally more similar to each other than Alpha, Beta and Delta^127^.

We observed many prominent shifts from complex glycans on Wuhan-Hu-1 to high mannose glycans on “variant” spikes, at multiple NGS **(Figure 3B)**. This trend hints at reduced *N*-glycan processivity at these NGS, hence implying an increase in their steric occlusion. It also suggests that variant spikes may be better at using mannose-binding membrane proteins as receptors for host-cell entry/fusion^131^, thus explaining SARS-CoV-2 tropism in the lower human respiratory tract, where infection is most prominent but the primary host receptor (ACE2) is minimally expressed^97,132^. This spike-glycan-based attachment of infectious virus to host cells can increase virus residence time on host surface, thus providing a better chance of host-virus fusion and consequent infection. Unlike the spike on an infectious virus, vaccine action is independent of cellular attachment or viral fusion, so the variant-based protein-structure changes are more likely to explain immunogenicity than the changes they effect in glycan landscapes. This notion was supported by studies that reported broader immunogenicity of partially-deglycosylated immunogens^133-135^. We benchmarked the Novavax SARS-CoV-2 spike and found its glycan landscape to be very similar to the 2P version of the Wuhan-Hu-1 spike (except significantly higher complex glycans at N801 and lower glycan occupancy at N657) **(Figure 5A,B)**, despite the absence of the TM and CT domains in the 2P version and both trimers being expressed in different cell types (2P version in human cells and Novavax vaccine in insect cells^59^). We note that this site-specific *N*-glycosylation pattern observed on Novavax SARS-CoV-2 spike-protein recapitulates very well the results from our previous attempt^59^ at analyzing this glycoprotein (the only substantial differences being at N17, N343, N657 and N717, which can be explained by scant sampling or high variance in our previous attempt), that was performed with an earlier version of DeGlyPHER^49,78^.

**Figure 5:**
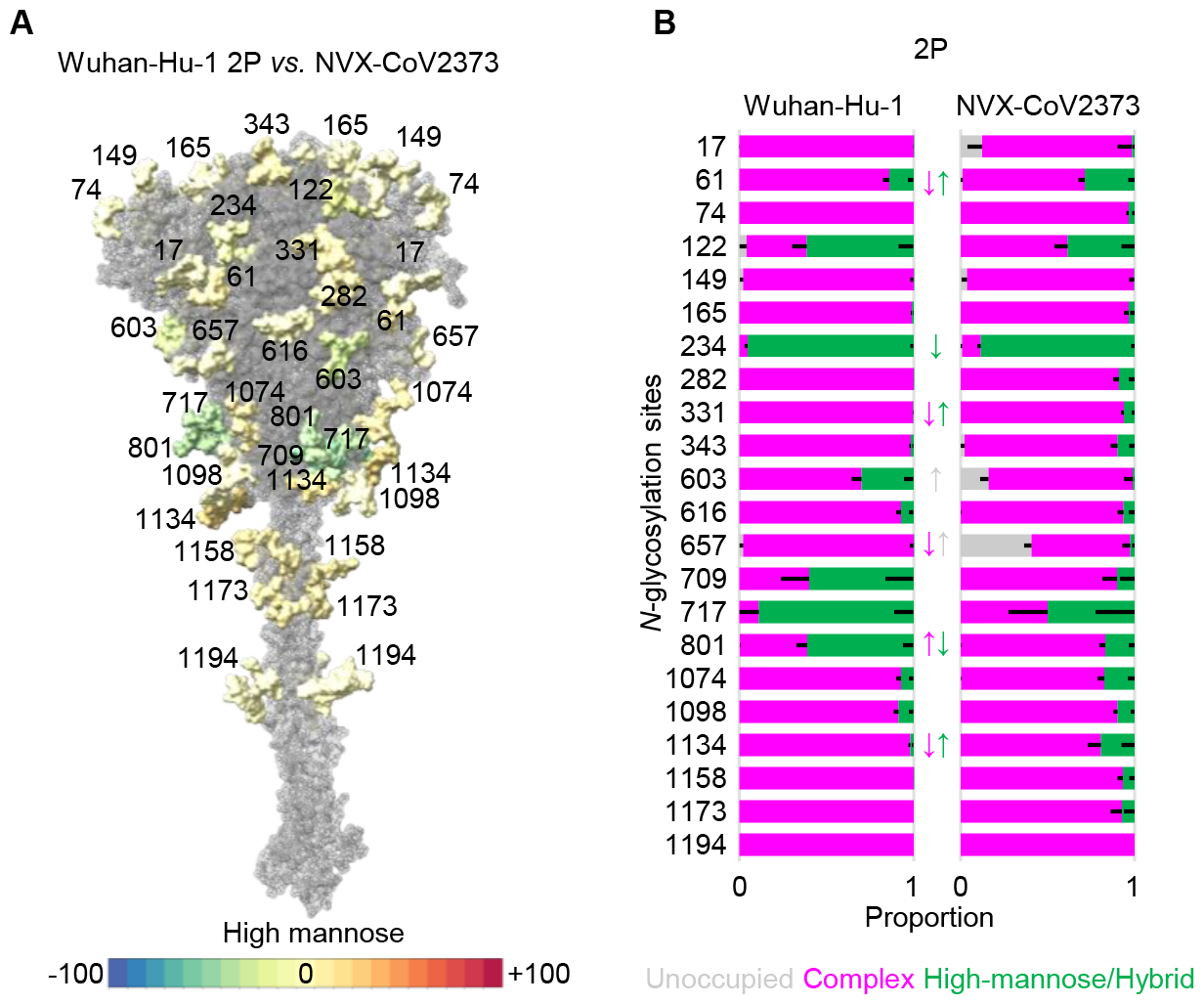
Comparing glycosylation changes between Wuhan-Hu-1 2P spike-protein and NVX-CoV2373. A percentage-point scale signifying increase (towards red) or decrease (towards blue) in high mannose *N*-glycans, shows not much difference between the 2 spike-protein trimers (A), even though many of the smaller differences are significant (B) owing much higher sampling from NVX-CoV2373. *N*-glycosylation states are color-coded. Error bars represent mean–SEM. Any significant difference (BH-corrected p-value <0.05) in proportion of a certain *N*-glycosylation state at an NGS is represented by color-coded ↕.

We observed the most conspicuous and consistent shifts from complex to high mannose glycans at N165, N343 and N616 **(Figure 3B)**. Newby *et. al*. see similar increase in high-mannose (mostly M5) glycans at N165 and N616, and hybrid glycans at N616^64^, and though not substantial, but Shajahan *et. al*.^65^ and Wang *et. al*.^66^ observe similar trends. Owing to structural changes in variant spike-proteins, Golgi α-mannosidase II (∼130 kDa), that initiates processing of all complex glycans^136^, may have limited access to process glycans at these NGS by virtue of its size, when compared to other mannosidases^137,138^ and glycosyl transferases^139,140^ (∼40-70 kDa) that form M4-M6 high-mannose and hybrid glycans^50^. N165 and N343 are implicated in RBD opening^52-54^, and N616 is a single amino acid away from D614G, which is the most consistent mutation in SARS-CoV-2 variants^9^ and implicated in critical structural changes that determine S1-S2 integrity after Furin cleavage^122,141^. So, there is very high probability that N616 glycans can regulate the integrity of S1-S2 coupling in the spike-protein trimer^85^. Presence of *O*-glycans has been reported at N618, that may also contribute to this effect^142,143^. P681R mutation may enhance Furin cleavage by adding a basic residue to 682RRAR in Delta and D614G stabilizes the Furin-cleaved S1-protein on S2-protein to facilitate fusogenicity, thus explaining Delta’s high disease severity^122,125,144,145^. The significant decrease in glycan processing at N616, N165 and N343 that we observed in Delta **(Figures 3A;4A,C)** may further explain fusogenicity and syncytia formation *via* mannose-binding receptors on respiratory tract cells that are the primary tissue affected by Delta but which sparsely express ACE2^93,97,110,131,146,147^. Further interrogation of this possibility may bring a critical advance in understanding Delta’s severe pathogenicity in human respiratory tissue.

SARS-CoV-2 spike is one of the most perused glycoprotein structures and a huge amount of structure-based immunology has been published on this since the COVID-19 pandemic^80,85,148-152^. Though these numerous studies often acknowledge the indispensable role of glycans and propose molecular mechanisms involving glycans that could possibly explain their observations and inferences, but there have been few studies evaluating site-specific glycosylation on spike-proteins from SARS-CoV-2 variants^64-66,153^. These MS-based traditional glycoproteomic approaches have provided insights into various glycoforms present at each site on spike-protein from SARS-CoV-2 variants, but their estimation of glycan occupancy and proportion of *N*-glycan types lacks precision, as evident from variation in estimation between studies and their inability to plot any metric of error to represent confidence in estimating proportion of glycan types they observe^57,60,64-66,70,76,135^; presumably due to inherent challenges of measuring sufficient glycopeptides mapping to each site, thus failing to recapitulate the accurate site-specific representative distribution of glycans with high degree of statistical confidence. This also limits their scope to make head-to-head or pairwise site-specific statistical comparison between variants. DeGlyPHER^77,79^ overcomes these limitations with excellent sampling per glycosylation site, providing the statistical power to make precise estimations and confident comparisons, and in rare cases where sampling dwindles, we abstain from making inferences. Thus, we used DeGlyPHER to compare the various VoC/Is of SARS-CoV-2 for their spike-protein *N*-glycosylations. DeGlyPHER’s advantage is evident from the large number of statistically confident glycosylation state differences we identified, and their critical importance is validated by many structural and immunological studies that are cited throughout this report.

## Acknowledgements

We thank Claire Delahunty and Sandhya Bangaru for critically reading the manuscript, Charles Bowman for computational assistance, Salvador Martínez-Bartolomé for assistance with GlycoMSQuant, and Alessio D’Addabbo and Max Crispin (University of Southampton, UK) for pointing us to the use of percentage-point scale. We thank Nita Patel, Alyse D. Portnoff, and Gale E. Smith from Novavax, Inc. for supplying NVX-CoV2373 for this research. This work was supported by the grants P41GM103533 (NIH), UM1AI100663 (NIH, CHAVI-ID), UM1AI144462 (NIH, CHAVD), R01/R56AI113867 (NIH/NIAID) and Bill and Melinda Gates Foundation (INV-004923).

## Contributions

S.B. conceptualized the project, performed glycan analysis, and wrote the manuscript with the contribution of all authors. J.K.D. performed LC-MS/MS. J.L.T and J.C. produced the spike-proteins. J.L.T performed the NS-EM. B.S. created the molecular structure graphics. P.T.G. performed the hierarchical clustering analysis. A.B.W., J.C.P, and J.R.Y supervised the project.

## Competing Interests

The authors declare no competing financial interest.

## Methods

### Glycoprotein expression and Purification

Spike-proteins (amino acid sequences in **Supplementary Information**) were expressed and purified as previously described^84^. Briefly, 500 μg of each plasmid encoding spike-proteins used in this study were incubated with 1.5 mg of polyethylenimine (Polysciences, Inc.) for 20 min and the mixture was deposited in 1L of HEK293F cells. Protein was expressed for 6 days, cells were harvested, and the protein was purified with a StrepTactin XT 4FLOW column (IBA Biosciences). The eluted protein was then size exclusion purified over a Superose 6 HiLoad 16/600 pg 120 mL or Superose 6 Increase 10/300 GL columns (Cytiva).

### Electron microscopy

To assess trimer homogeneity, 3 μL of spike-protein at 0.03 mg/ml in 1X TBS was deposited on a glow-discharged carbon coated copper mesh grid (Electron Microscopy Sciences) and blotted off. 3 μL of 2% uranyl formate (Electron Microscopy Sciences) was deposited on the grid for 90 s then blotted off. Data collection was done on a FEI Tecnai Spirit (120 keV) and a FEI Eagle 4k x 4k CCD camera (Thermo Fisher) and automated with the Leginon software^154^. Raw micrographs were stored in the Appion^155^ database, and 2D classes were generated with RELION^156^.

### Mass spectrometry analysis

DeGlyPHER^77,79^ was used to ascertain site-specific glycan occupancy and processivity on glycoproteins.

#### Proteinase K treatment and deglycosylation

Purified SARS-CoV-2 spike glycoproteins were exchanged to water using Microcon Ultracel PL-10 centrifugal filter (Millipore Sigma). Glycoproteins were reduced with 5 mM tris(2-carboxyethyl)phosphine hydrochloride (TCEP-HCl; Thermo Scientific) and alkylated with 10 mM 2-Chloroacetamide (Sigma-Aldrich) in 100 mM ammonium acetate (Sigma Aldrich) for 20 min at room temperature (RT, 24LC). Initial protein-level deglycosylation was performed using 250 U of Endo H (New England Biolabs) for 5 µg trimer, for 1 h at 37□C. Glycoproteins were digested with 1:25 Proteinase K (PK; Sigma-Aldrich) for 30 min at 37□C (For NVX-CoV2373, 9 different conditions were used *viz*., 16 h of trypsin (Promega) digestion, and variations in PK treatment – 1:25 PK for 30 min, 1 h, 2 h, 4 h; 1:100, 1:500, 1:1000, 1:2000 PK for 4 h). PK was denatured by incubating at 90□C for 15 min, then cooled to RT. Peptides were deglycosylated again with 250 U Endo H for 1 h at 37□C, then frozen at –80□C and lyophilized. 100 U PNGase F (New England Biolabs) was lyophilized, resuspended in 20 µl 100 mM ammonium bicarbonate (Sigma-Aldrich) prepared in H_2_ ^18^O (Sigma-Aldrich), and added to the lyophilized peptides. Reactions were then incubated for 1 h at 37_□_C, and subsequently analyzed by LC-MS/MS.

#### LC-MS/MS

Samples were analyzed on an Q Exactive HF-X mass spectrometer (Thermo Scientific). Samples were injected directly onto a 25 cm, 100 μm ID column packed with BEH 1.7 μm C18 resin (Waters). Samples were separated at a flow rate of 300 nL/min on an EASY-nLC 1200 UHPLC (Thermo Scientific). Buffers A and B were 0.1% formic acid in 5% and 80% acetonitrile, respectively. The following gradient was used: 1–25% B over 160 min, an increase to 40% B over 40 min, an increase to 90% B over another 10 min and 30 min at 90% B for a total run time of 240 min. Column was re-equilibrated with Buffer A prior to the injection of sample. Peptides were eluted from the tip of the column and nanosprayed directly into the mass spectrometer by application of 2.8 kV at the back of the column. The mass spectrometer was operated in a data dependent mode. Full MS1 scans were collected in the Orbitrap at 120,000 resolution. The ten most abundant ions per scan were selected for HCD MS/MS at 25 NCE. Dynamic exclusion was enabled with exclusion duration of 10 s and singly charged ions were excluded.

#### Data Processing

Protein and peptide identifications were made with Integrated Proteomics Pipeline (IP2). Tandem mass spectra were extracted from raw files using RawConverter^157^ and searched with ProLuCID^158^ against a database comprising UniProt reviewed (Swiss-Prot) proteome for Homo sapiens (UP000005640), UniProt amino acid sequences for Endo H (P04067), PNGase F (Q9XBM8), and Proteinase K (P06873), amino acid sequences for the examined proteins, and a list of general protein contaminants. The search space included no cleavage-specificity. Carbamidomethylation (+57.02146 C) was considered a static modification. Deamidation in presence of H_2_ ^18^O (+2.988261 N), GlcNAc (+203.079373 N), oxidation (+15.994915 M) and N-terminal pyroglutamate formation (–17.026549 Q) were considered differential modifications. Data was searched with 50 ppm precursor ion tolerance and 50 ppm fragment ion tolerance. Identified proteins were filtered using DTASelect2^159^, utilizing a target-decoy database search strategy to limit the false discovery rate to 1%, at the spectrum level^160^. A minimum of 1 peptide per protein and no tryptic end per peptide were required and precursor delta mass cut-off was fixed at 15 ppm. Statistical models for peptide mass modification (modstat) were applied. Census2^161^ label-free analysis was performed based on the precursor peak area, with a 15 ppm precursor mass tolerance and 0.1 min retention time tolerance. “Match between runs” was used to find missing peptides between runs. Data analysis using GlycoMSQuant^77^ was implemented to automate the analysis. GlycoMSQuant summed precursor peak areas across replicates, discarded peptides without NGS, discarded misidentified peptides when *N*-glycan remnant-mass modifications were localized to non-NGS asparagines and corrected/fixed *N*-glycan mislocalization where appropriate. The results were aligned to NGS in spike of Wuhan-Hu-1 (UniProt #P0DTC2). The data can be accessed from MassIVE (MSV000091921)

A glycosylated model^53^ based on the “closed state” of SARS-CoV-2 spike-protein trimer (PDB: 6VXX)^42^ and *N*-glycosylation profile determined by Watanabe *et. al*.^57^ was uploaded on ChimeraX v.1.5 (https://www.cgl.ucsf.edu/chimerax/). Glycans were separated into separate PDB files using The PyMOL Molecular Graphics System v.2.5.2 (https://pymol.org/). Coordinates for membrane structure were hidden and glycans were color-coded using Python v.3.9.7 (https://www.python.org/) in multiple formats: [1] monochrome to highlight all glycans over protein sub-structure, [2] a 5-color gradient to classify the dominant proportion of unoccupied glycosylation sites, high-mannose/hybrid glycans and complex glycans, and [3] a 21-color gradient to represent percentage-point shift in proportion of high-mannose/hybrid glycans at a particular glycosylation site^162^, when comparing 2 immunogens. The code used to create visualizations can be accessed from GitHub (https://github.com/BhavyaSingh-Rolly/SARS-Visualization)

Proportional distribution of glycans at NGS common to all variants (*i.e*., except N17, N20, N74 and N188, and the inconsistently sampled N709, N717 and N1134), as calculated by GlycoMSQuant^77^, was condensed into a 1D feature vector, and a pairwise correlation analysis was conducted between all variants in their 6P-Mut7 versions. Subsequently, we used Scipy’s hierarchical linkage function to cluster based on similarity and then used the hierarchical dendrogram function to visualize the results^163^.

## Supplementary Information Titles/Legends

**Figure S1: Previous quantitation of site-specific *N*-glycosylation on SARS-CoV-2 spike-protein trimer vary**. Shown are the results from site-specific *N*-glycosylation analysis by standard MS-based glycoproteomic method on Wuhan-Hu-1 as reported by Watanabe *et. al*., 2020 [S1] (Report 1); Chawla *et. al*., 2022 [S2] (Report 2); Newby *et. al*., 2023 [S3] (Report 3); Shajahan *et. al*., 2020 [S4] (Report 4); Shajahan *et. al*., 2023 [S5] (Report 5); and Wang *et. al*., 2021 [S6] (Report 6), compared to results from DeGlyPHER (top 2 rows, 2P followed by 6P). A visual estimate from these reports is plotted here and compared at each *N*-glycosylation site. Variation in glycosylation pattern observed in different reports are apparent at many sites. Color-coding of “Report #”, groups the reports based on research laboratories authoring them. Cited studies used either 2P stabilized mutant of SARS-CoV-2 spike-protein [S7, S8] or 6P stabilized mutant [S9], as indicated, except report 4 that used independently expressed S1 and S2 subunits of S-protein. Error bars are absent here because all the cited reports here except this study, have not estimated error in their calculations, presumably owing to low sampling. *Report 4 does not claim quantitation; thus, estimated values are from the types of glycoforms reported. Only this study and Report 1 have validated that S-proteins examined are well-folded trimers (using negative-stain electron microscopy). n.d.: glycosylation not determined.

**Figure S2: Evaluating SARS-CoV-2 spike-protein trimer purity and integrity**. (A) Size exclusion chromatography of SARS-CoV-2 spike-protein trimers on Superose 6 columns. Trimer elution peaks at 68-and 13-mL post-injection with Superose 6 Hiload 16/600 pg and Superose 6 Increase 10/300 GL columns, respectively. (B) Representative negative-stain TEM micrographs and 2D classes of SARS-CoV-2 spike-protein trimers show highly homogenous particles in various orientations.

**Figure S3: Peptide sampling at each NGS across all spike-protein trimers analyzed**. (A) Number of unique peptides mapping to each NGS shows that N709, N717 and N1134 are inconsistently sampled, but all other NGS are sufficiently sampled. (B) Bar graph demonstrating *N*-glycan heterogeneity determined at N1134. The *N*-glycans present at N1134 are mostly complex as seen in previous studies, though the sampling at this NGS is inconsistent in our study. *N*-glycosylation states are color-coded. Error bars represent mean–SEM.

**Supplementary Table-1: SARS-CoV-2 variant spike proteins**.

**Supplementary Table-2: SARS-CoV-2 spike-protein stabilizing mutations**.

**SARS-CoV-2 spike-protein sequences**.

## Notes

### Competing Interest Statement

The authors have declared no competing interest.

### Summary of Updates

New updated supplemental file that includes a new figure (S1) to help discuss the results. New references added.

