## Supporting Information for "Evolving spike-protein *N*-glycosylation in SARS-CoV-2 variants"

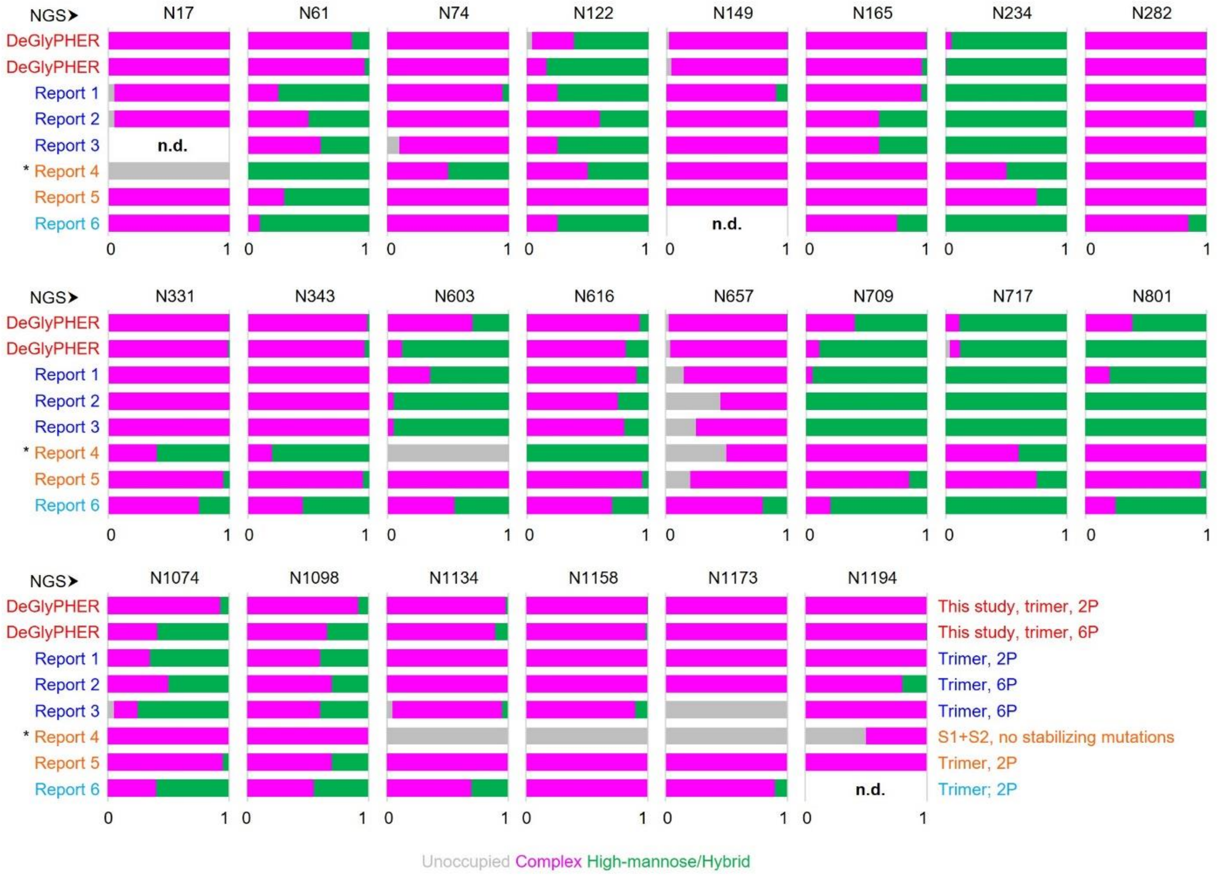


**Figure S1: Previous quantitation of site-specific N-glycosylation on SARS-CoV-2 spike-protein trimer vary.** Shown are the results from site-specific N-glycosylation analysis by standard MS-based glycoproteomic method on Wuhan-Hu-1 as reported by Watanabe et. al., 2020 [S1] (Report 1); Chawla et. al., 2022 [S2] (Report 2); Newby et. al., 2023 [S3] (Report 3); Shajahan et. al., 2020 [S4] (Report 4); Shajahan et. al., 2023 [S5] (Report 5); and Wang et. al., 2021 [S6] (Report 6), compared to results from DeGlyPHER (top 2 rows, 2P followed by 6P). A visual estimate from these reports is plotted here and compared at each N-glycosylation site. Variation in glycosylation pattern observed in different reports are apparent at many sites. Color-coding of "Report #", groups the reports based on research laboratories authoring them. Cited studies used either 2P stabilized mutant of SARS-CoV-2 spike-protein [S7, S8] or 6P stabilized mutant [S9], as indicated, except report 4 that used independently expressed S1 and S2 subunits of S-protein. Error bars are absent here because all the cited reports here except this study, have not estimated error in their calculations, presumably owing to low sampling. *Report 4 does not claim quantitation; thus, estimated values are from the types of glycoforms reported. Only this study and Report 1 have validated that S-proteins examined are well-folded trimers (using negative-stain electron microscopy). n.d.: glycosylation not determined.

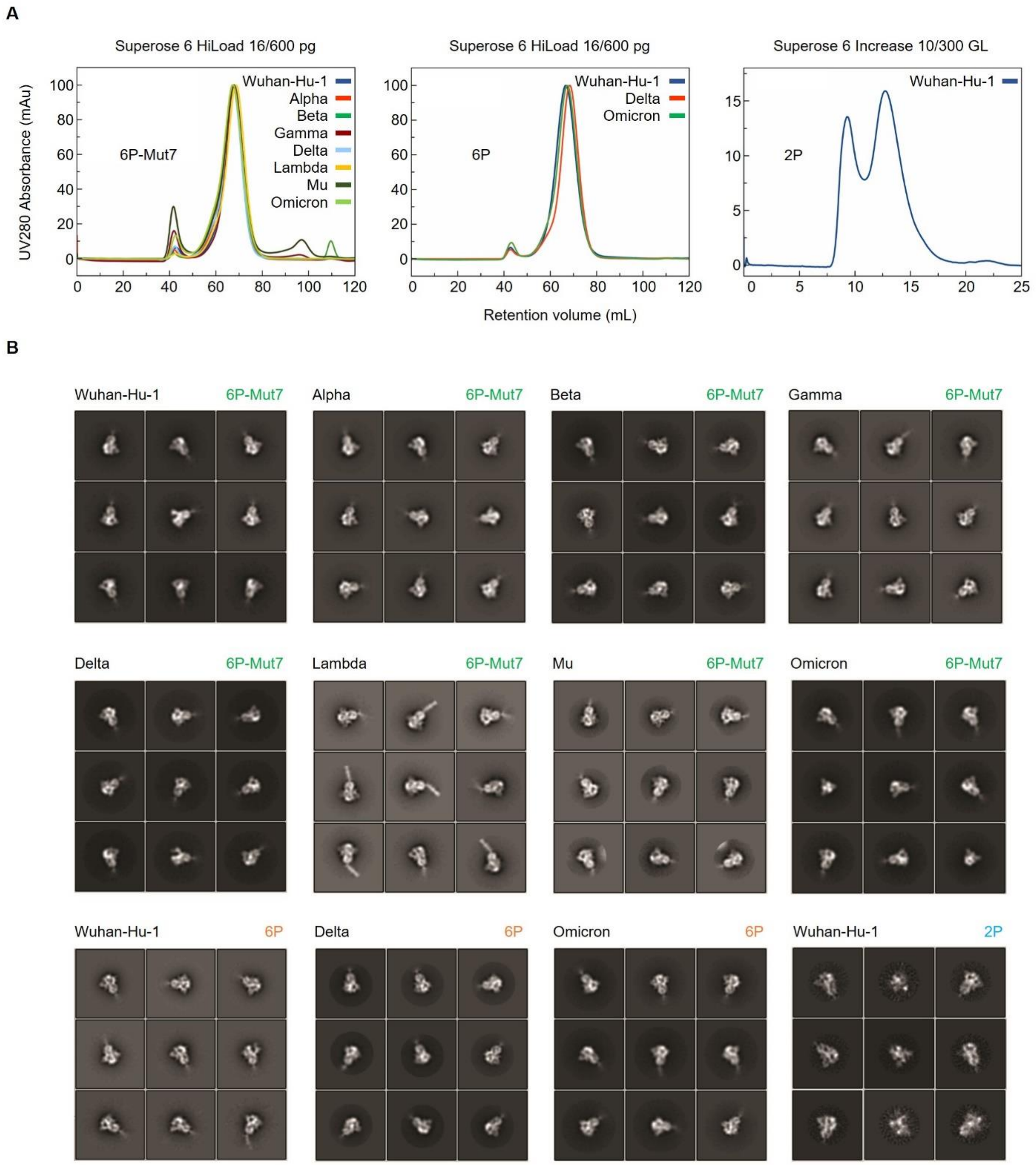


**Figure S2: Evaluating SARS-CoV-2 spike-protein trimer purity and integrity.** (A) Size exclusion chromatography of SARS-CoV-2 spike-protein trimers on Superose 6 columns. Trimer elution peaks at 68- and 13-mL post-injection with Superose 6 Hiload 16/600 pg and Superose 6 Increase 10/300 GL columns, respectively. (B) Representative negative-stain TEM micrographs and 2D classes of SARS-CoV-2 spike-protein trimers show highly homogenous particles in various orientations.

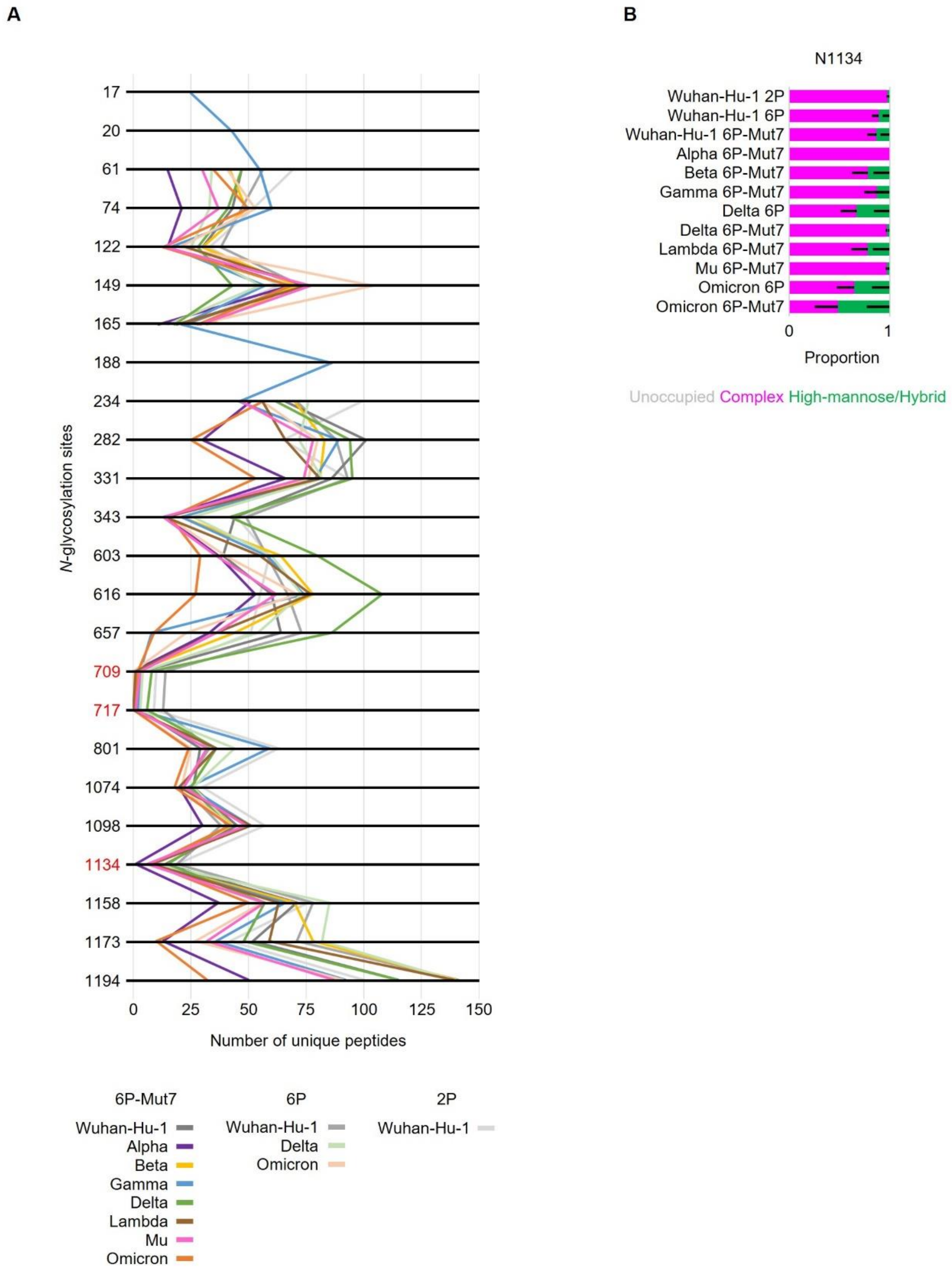


**Figure S3: Peptide sampling at each NGS across all spike-protein trimers analyzed.** (A) Number of unique peptides mapping to each NGS shows that N709, N717 and N1134 are inconsistently sampled, but all other NGS are sufficiently sampled. (B) Bar graph demonstrating N-glycan heterogeneity determined at N1134. The *N*-glycans present at N1134 are mostly complex as seen in previous studies, though the sampling at this NGS is inconsistent in our study. *N*-glycosylation states are color-coded. Error bars represent mean–SEM.

**Supplementary Table-1: SARS-CoV-2 variant spike proteins.**

| Variants | Pango | Mutations | Stabilization | Length | Furin |
|---|---|---|---|---|---|
| **Wuhan-Hu-1** | A | - | 2P, 6P, 6P-Mut7 | 1-1208 | GSAS |
| **Alpha** | B.1.1.7 | del69-70, del144, N501Y, A570D, D614G, P681H, T716I, S982A, D1118H | 6P-Mut7 | 1-1208 | GSAS |
| **Beta** | B.1.351 | L18F, D80A, D215G, del242-244, K417N, E484K, N501Y, D614G, A701V | 6P-Mut7 | 1-1208 | GSAS |
| **Gamma** | P.1 | L18F, T20N, P26S, D138Y, R190S, K417T, E484K, N501Y, D614G, H655Y, T1027I, V1176F | 6P-Mut7 | 1-1208 | GSAS |
| **Delta** | B.1.617.2 | T19R, G142D, del156-157, R158G, L452R, T478K, D614G, P681R, D950N | 6P, 6P-Mut7 | 1-1208 | GSAS |
| **Omicron** | B.1.1.529 | A67V, del69-70, T95I, G142D, del143-145, del211, ins214EPE, G339D, S371L, S373P, S375F, K417N, N440K, G446S, S477N, T478K, E484A, Q493R, G496S, Q498R, N501Y, Y505H, T547K, D614G, H655Y, N679K, P681H, N764K, D796Y, N856K, Q954H, N969K, L981F | 6P, 6P-Mut7 | 1-1208 | GSAS |
| **Mu** | B1.1.621 | T95I, Y144T, Y145S, ins146N, R346K, E484K, N501Y, D614G, P681H, D950N | 6P-Mut7 | 1-1208 | GSAS |
| **Lambda** | C.37 | G75V, T76I, del246-252, L452Q, F490S, D614G, T859N | 6P-Mut7 | 1-1208 | GSAS |
| **NVX-CoV2373** | - | - | 2P | 1-1273 | QQAQ |

**Supplementary Table-2: SARS-CoV-2 spike-protein stabilizing mutations.**

| Versions | Mutations |
|---|---|
| **2P** | K986P, V987P |
| **6P** | F817P, A892P, A899P, A942P, K986P, V987P |
| **6P-Mut7** | F817P, A892P, A899P, A942P, K986P, V987P; **Mut7** – V705C, T883C |

**SARS-CoV-2 spike-protein sequences.**

**Wuhan-Hu-1 2P, 1-1208 (2P – K986P, V987P; Furin CS – R682G, R683S, R685S)**

MFVFLVLLPLVSSQCVNLTTRTQLPPAYTNSFTRGVYYPDKVFRSSVLHSTQDLFLPFFSNVTWFHAIHV
SGTNGTKRFDNPVLPFNDGVYFASTEKSNIIRGWIFGTTLDSKTQSLLIVNNATNVVIKVCEFQFCNDPF
LGVYYHKNNKSWMESEFRVYSSANNCTFEYVSQPFLMDLEGKQGNFKNLREFVFKNIDGYFKIYSKHTPI
NLVRDLPQGFSALEPLVDLPIGINITRFQTLLALHRSYLTPGDSSSGWTAGAAAYYVGYLQPRTFLLKYN
ENGTITDAVDCALDPLSETKCTLKSFTVEKGIYQTSNFRVQPTESIVRFPNITNLCPFGEVFNATRFASV
YAWNRKRISNCVADYSVLYNSASFSTFKCYGVSPTKLNDLCFTNVYADSFVIRGDEVRQIAPGQTGKIAD
YNYKLPDDFTGCVIAWNSNNLDSKVGGNYNYLYRLFRKSNLKPFERDISTEIYQAGSTPCNGVEGFNCYF
PLQSYGFQPTNGVGYQPYRVVVLSFELLHAPATVCGPKKSTNLVKNKCVNFNFNGLTGTGVLTESNKKFL
PFQQFGRDIADTTDAVRDPQTLEILDITPCSFGGVSVITPGTNTSNQVAVLYQDVNCTEVPVAIHADQLT
PTWRVYSTGSNVFQTRAGCLIGAEHVNNSYECDIPIGAGICASYQTQTNSPGSASSVASQSIIAYTMSLG
AENSVAYSNNSIAIPTNFTISVTTEILPVSMTKTSVDCTMYICGDSTECSNLLLQYGSFCTQLNRALTGI
AVEQDKNTQEVFAQVKQIYKTPPIKDFGGFNFSQILPDPSKPSKRSFIEDLLFNKVTLADAGFIKQYGDC
LGDIAARDLICAQKFNGLTVLPPLLTDEMIAQYTSALLAGTITSGWTFGAGAALQIPFAMQMAYRFNGIG
VTQNVLYENQKLIANQFNSAIGKIQDSLSSTASALGKLQDVVNQNAQALNTLVKQLSSNFGAISSVLNDI
LSRLDPPEAEVQIDRLITGRLQSLQTYVTQQLIRAAEIRASANLAATKMSECVLGQSKRVDFCGKGYHLM
SFPQSAPHGVVFLHVTYVPAQEKNFTTAPAICHDGKAHFPREGVFVSNGTHWFVTQRNFYEPQIITTDNT
FVSGNCDVVIGIVNNTVYDPLQPELDSFKEELDKYFKNHTSPDVDLGDISGINASVVNIQKEIDRLNEVA
KNLNESLIDLQELGKYEQGSGYIPEAPRDGQAYVRKDGEWVLLSTFLGRSLEVLFQGPGHHHHHHHHSAW
SHPQFEKGGGSGGGGSGGSAWSHPQFEK*

**Wuhan-Hu-1 6P, 1-1208 (6P – F817P, A892P, A899P, A942P, K986P, V987P; Furin CS – R682G, R683S, R685S)**

MFVFLVLLPLVSSQCVNLTTRTQLPPAYTNSFTRGVYYPDKVFRSSVLHSTQDLFLPFFSNVTWFHAIHV
SGTNGTKRFDNPVLPFNDGVYFASTEKSNIIRGWIFGTTLDSKTQSLLIVNNATNVVIKVCEFQFCNDPF
LGVYYHKNNKSWMESEFRVYSSANNCTFEYVSQPFLMDLEGKQGNFKNLREFVFKNIDGYFKIYSKHTPI
NLVRDLPQGFSALEPLVDLPIGINITRFQTLLALHRSYLTPGDSSSGWTAGAAAYYVGYLQPRTFLLKYN
ENGTITDAVDCALDPLSETKCTLKSFTVEKGIYQTSNFRVQPTESIVRFPNITNLCPFGEVFNATRFASV
YAWNRKRISNCVADYSVLYNSASFSTFKCYGVSPTKLNDLCFTNVYADSFVIRGDEVRQIAPGQTGKIAD
YNYKLPDDFTGCVIAWNSNNLDSKVGGNYNYLYRLFRKSNLKPFERDISTEIYQAGSTPCNGVEGFNCYF
PLQSYGFQPTNGVGYQPYRVVVLSFELLHAPATVCGPKKSTNLVKNKCVNFNFNGLTGTGVLTESNKKFL
PFQQFGRDIADTTDAVRDPQTLEILDITPCSFGGVSVITPGTNTSNQVAVLYQDVNCTEVPVAIHADQLT
PTWRVYSTGSNVFQTRAGCLIGAEHVNNSYECDIPIGAGICASYQTQTNSPGSASSVASQSIIAYTMSLG
AENSVAYSNNSIAIPTNFTISVTTEILPVSMTKTSVDCTMYICGDSTECSNLLLQYGSFCTQLNRALTGI
AVEQDKNTQEVFAQVKQIYKTPPIKDFGGFNFSQILPDPSKPSKRSPIEDLLFNKVTLADAGFIKQYGDC
LGDIAARDLICAQKFNGLTVLPPLLTDEMIAQYTSALLAGTITSGWTFGAGPALQIPFPMQMAYRFNGIG
VTQNVLYENQKLIANQFNSAIGKIQDSLSSTPSALGKLQDVVNQNAQALNTLVKQLSSNFGAISSVLNDI
LSRLDPPEAEVQIDRLITGRLQSLQTYVTQQLIRAAEIRASANLAATKMSECVLGQSKRVDFCGKGYHLM
SFPQSAPHGVVFLHVTYVPAQEKNFTTAPAICHDGKAHFPREGVFVSNGTHWFVTQRNFYEPQIITTDNT
FVSGNCDVVIGIVNNTVYDPLQPELDSFKEELDKYFKNHTSPDVDLGDISGINASVVNIQKEIDRLNEVA
KNLNESLIDLQELGKYEQGSGYIPEAPRDGQAYVRKDGEWVLLSTFLGRSLEVLFQGPGSAWSHPQFEKG
GGSGGGGSGGSAWSHPQFEK*

**Wuhan-Hu-1 6P-Mut7, 1-1208 (6P – F817P, A892P, A899P, A942P, K986P, V987P; Mut7 – V705C, T883C; Furin CS – R682G, R683S, R685S)**

MFVFLVLLPLVSSQCVNLTTRTQLPPAYTNSFTRGVYYPDKVFRSSVLHSTQDLFLPFFSNVTWFHAIHV
SGTNGTKRFDNPVLPFNDGVYFASTEKSNIIRGWIFGTTLDSKTQSLLIVNNATNVVIKVCEFQFCNDPF
LGVYYHKNNKSWMESEFRVYSSANNCTFEYVSQPFLMDLEGKQGNFKNLREFVFKNIDGYFKIYSKHTPI
NLVRDLPQGFSALEPLVDLPIGINITRFQTLLALHRSYLTPGDSSSGWTAGAAAYYVGYLQPRTFLLKYN
ENGTITDAVDCALDPLSETKCTLKSFTVEKGIYQTSNFRVQPTESIVRFPNITNLCPFGEVFNATRFASV

YAWNRKRISNCVADYSVLYNSASFSTFKCYGVSPTKLNDLCFTNVYADSFVIRGDEVRQIAPGQTGKIAD
YNYKLPDDFTGCVIAWNSNNLDSKVGGNYNYLYRLFRKSNLKPFERDISTEIYQAGSTPCNGVEGFNCYF
PLQSYGFQPTNGVGYQPYRVVVLSFELLHAPATVCGPKKSTNLVKNKCVNFNFNGLTGTGVLTESNKKFL
PFQQFGRDIADTTDAVRDPQTLEILDITPCSFGGVSVITPGTNTSNQVAVLYQDVNCTEVPVAIHADQLT
PTWRVYSTGSNVFQTRAGCLIGAEHVNNSYECDIPIGAGICASYQTQTNSPGSASSVASQSIIAYTMSLG
AENSCAYSNNSIAIPTNFTISVTTEILPVSMTKTSVDCTMYICGDSTECSNLLLQYGSFCTQLNRALTGI
AVEQDKNTQEVFAQVKQIYKTPPIKDFGGFNFSQILPDPSKPSKRSPIEDLLFNKVTLADAGFIKQYGDC
LGDIAARDLICAQKFNGLTVLPPLLTDEMIAQYTSALLAGTICSGWTFGAGPALQIPFPMQMAYRFNGIG
VTQNVLYENQKLIANQFNSAIGKIQDSLSSTPSALGKLQDVVNQNAQALNTLVKQLSSNFGAISSVLNDI
LSRLDPPEAEVQIDRLITGRLQSLQTYVTQQLIRAAEIRASANLAATKMSECVLGQSKRVDFCGKGYHLM
SFPQSAPHGVVFLHVTYVPAQEKNFTTAPAICHDGKAHFPREGVFVSNGTHWFVTQRNFYEPQIITTDNT
FVSGNCDVVIGIVNNTVYDPLQPELDSFKEELDKYFKNHTSPDVDLGDISGINASVVNIQKEIDRLNEVA
KNLNESLIDLQELGKYEQGSGYIPEAPRDGQAYVRKDGEWVLLSTFLGRSLEVLFQGPGSAWSHPQFEKG
GGSGGGGSGGSAWSHPQFEK*

**Alpha (B.1.1.7) Variant 6P-Mut7, 1-1208 (del69-70, del144, N501Y, A570D, D614G, P681H, T716I, S982A, D1118H; 6P – F817P, A892P, A899P, A942P, K986P, V987P; Mut7 – V705C, T883C; Furin CS – R682G, R683S, R685S)**

MFVFLVLLPLVSSQCVNLTTRTQLPPAYTNSFTRGVYYPDKVFRSSVLHSTQDLFLPFFSNVTWFHAISG
TNGTKRFDNPVLPFNDGVYFASTEKSNIIRGWIFGTTLDSKTQSLLIVNNATNVVIKVCEFQFCNDPFLG
VYHKNNKSWMESEFRVYSSANNCTFEYVSQPFLMDLEGKQGNFKNLREFVFKNIDGYFKIYSKHTPINLV
RDLPQGFSALEPLVDLPIGINITRFQTLLALHRSYLTPGDSSSGWTAGAAAYYVGYLQPRTFLLKYNENG
TITDAVDCALDPLSETKCTLKSFTVEKGIYQTSNFRVQPTESIVRFPNITNLCPFGEVFNATRFASVYAW
NRKRISNCVADYSVLYNSASFSTFKCYGVSPTKLNDLCFTNVYADSFVIRGDEVRQIAPGQTGKIADYNY
KLPDDFTGCVIAWNSNNLDSKVGGNYNYLYRLFRKSNLKPFERDISTEIYQAGSTPCNGVEGFNCYFPLQ
SYGFQPTYGVGYQPYRVVVLSFELLHAPATVCGPKKSTNLVKNKCVNFNFNGLTGTGVLTESNKKFLPFQ
QFGRDIDDTTDAVRDPQTLEILDITPCSFGGVSVITPGTNTSNQVAVLYQGVNCTEVPVAIHADQLTPTW
RVYSTGSNVFQTRAGCLIGAEHVNNSYECDIPIGAGICASYQTQTNSHGSASSVASQSIIAYTMSLGAEN
SCAYSNNSIAIPINFTISVTTEILPVSMTKTSVDCTMYICGDSTECSNLLLQYGSFCTQLNRALTGIAVE
QDKNTQEVFAQVKQIYKTPPIKDFGGFNFSQILPDPSKPSKRSPIEDLLFNKVTLADAGFIKQYGDCLGD
IAARDLICAQKFNGLTVLPPLLTDEMIAQYTSALLAGTICSGWTFGAGPALQIPFPMQMAYRFNGIGVTQ
NVLYENQKLIANQFNSAIGKIQDSLSSTPSALGKLQDVVNQNAQALNTLVKQLSSNFGAISSVLNDILAR
LDPPEAEVQIDRLITGRLQSLQTYVTQQLIRAAEIRASANLAATKMSECVLGQSKRVDFCGKGYHLMSFP
QSAPHGVVFLHVTYVPAQEKNFTTAPAICHDGKAHFPREGVFVSNGTHWFVTQRNFYEPQIITTHNTFVS
GNCDVVIGIVNNTVYDPLQPELDSFKEELDKYFKNHTSPDVDLGDISGINASVVNIQKEIDRLNEVAKNL
NESLIDLQELGKYEQGSGYIPEAPRDGQAYVRKDGEWVLLSTFLGRSLEVLFQGPGSAWSHPQFEKGGGS
GGGGSGGSAWSHPQFEK*

**Beta (B.1.351) Variant 6P-Mut7, 1-1208 (L18F, D80A, D215G, del242-244, K417N, E484K, N501Y, D614G, A701V; 6P – F817P, A892P, A899P, A942P, K986P, V987P; Mut7 – V705C, T883C; Furin CS – R682G, R683S, R685S)**

MFVFLVLLPLVSSQCVNFTTRTQLPPAYTNSFTRGVYYPDKVFRSSVLHSTQDLFLPFFSNVTWFHAIHV
SGTNGTKRFANPVLPFNDGVYFASTEKSNIIRGWIFGTTLDSKTQSLLIVNNATNVVIKVCEFQFCNDPF
LGVYYHKNNKSWMESEFRVYSSANNCTFEYVSQPFLMDLEGKQGNFKNLREFVFKNIDGYFKIYSKHTPI
NLVRGLPQGFSALEPLVDLPIGINITRFQTLHRSYLTPGDSSSGWTAGAAAYYVGYLQPRTFLLKYNENG
TITDAVDCALDPLSETKCTLKSFTVEKGIYQTSNFRVQPTESIVRFPNITNLCPFGEVFNATRFASVYAW
NRKRISNCVADYSVLYNSASFSTFKCYGVSPTKLNDLCFTNVYADSFVIRGDEVRQIAPGQTGNIADYNY
KLPDDFTGCVIAWNSNNLDSKVGGNYNYLYRLFRKSNLKPFERDISTEIYQAGSTPCNGVKGFNCYFPLQ
SYGFQPTYGVGYQPYRVVVLSFELLHAPATVCGPKKSTNLVKNKCVNFNFNGLTGTGVLTESNKKFLPFQ
QFGRDIADTTDAVRDPQTLEILDITPCSFGGVSVITPGTNTSNQVAVLYQGVNCTEVPVAIHADQLTPTW
RVYSTGSNVFQTRAGCLIGAEHVNNSYECDIPIGAGICASYQTQTNSPGSASSVASQSIIAYTMSLGVEN
SCAYSNNSIAIPTNFTISVTTEILPVSMTKTSVDCTMYICGDSTECSNLLLQYGSFCTQLNRALTGIAVE

QDKNTQEVFAQVKQIYKTPPIKDFGGFNFSQILPDPSKPSKRSPIEDLLFNKVTLADAGFIKQYGDCLGD
IAARDLICAQKFNGLTVLPPLLTDEMIAQYTSALLAGTICSGWTFGAGPALQIPFPMQMAYRFNGIGVTQ
NVLYENQKLIANQFNSAIGKIQDSLSSTPSALGKLQDVVNQNAQALNTLVKQLSSNFGAISSVLNDILSR
LDPPEAEVQIDRLITGRLQSLQTYVTQQLIRAAEIRASANLAATKMSECVLGQSKRVDFCGKGYHLMSFP
QSAPHGVVFLHVTYVPAQEKNFTTAPAICHDGKAHFPREGVFVSNGTHWFVTQRNFYEPQIITTDNTFVS
GNCDVVIGIVNNTVYDPLQPELDSFKEELDKYFKNHTSPDVDLGDISGINASVVNIQKEIDRLNEVAKNL
NESLIDLQELGKYEQGSGYIPEAPRDGQAYVRKDGEWVLLSTFLGRSLEVLFQGPGSAWSHPQFEKGGGS
GGGGSGGSAWSHPQFEK*

**Gamma (P.1) Variant 6P-Mut7, 1-1208 (L18F, T20N, P26S, D138Y, R190S, K417T, E484K, N501Y, D614G, H655Y, T1027I, V1176F; 6P – F817P, A892P, A899P, A942P, K986P, V987P; Mut7 – V705C, T883C; Furin CS – R682G, R683S, R685S)**

MFVFLVLLPLVSSQCVNFTNRTQLPSAYTNSFTRGVYYPDKVFRSSVLHSTQDLFLPFFSNVTWFHAIHV
SGTNGTKRFDNPVLPFNDGVYFASTEKSNIIRGWIFGTTLDSKTQSLLIVNNATNVVIKVCEFQFCNYPF
LGVYYHKNNKSWMESEFRVYSSANNCTFEYVSQPFLMDLEGKQGNFKNLSEFVFKNIDGYFKIYSKHTPI
NLVRDLPQGFSALEPLVDLPIGINITRFQTLLALHRSYLTPGDSSSGWTAGAAAYYVGYLQPRTFLLKYN
ENGTITDAVDCALDPLSETKCTLKSFTVEKGIYQTSNFRVQPTESIVRFPNITNLCPFGEVFNATRFASV
YAWNRKRISNCVADYSVLYNSASFSTFKCYGVSPTKLNDLCFTNVYADSFVIRGDEVRQIAPGQTGTIAD
YNYKLPDDFTGCVIAWNSNNLDSKVGGNYNYLYRLFRKSNLKPFERDISTEIYQAGSTPCNGVKGFNCYF
PLQSYGFQPTYGVGYQPYRVVVLSFELLHAPATVCGPKKSTNLVKNKCVNFNFNGLTGTGVLTESNKKFL
PFQQFGRDIADTTDAVRDPQTLEILDITPCSFGGVSVITPGTNTSNQVAVLYQGVNCTEVPVAIHADQLT
PTWRVYSTGSNVFQTRAGCLIGAEYVNNSYECDIPIGAGICASYQTQTNSPGSASSVASQSIIAYTMSLG
AENSCAYSNNSIAIPTNFTISVTTEILPVSMTKTSVDCTMYICGDSTECSNLLLQYGSFCTQLNRALTGI
AVEQDKNTQEVFAQVKQIYKTPPIKDFGGFNFSQILPDPSKPSKRSPIEDLLFNKVTLADAGFIKQYGDC
LGDIAARDLICAQKFNGLTVLPPLLTDEMIAQYTSALLAGTICSGWTFGAGPALQIPFPMQMAYRFNGIG
VTQNVLYENQKLIANQFNSAIGKIQDSLSSTPSALGKLQDVVNQNAQALNTLVKQLSSNFGAISSVLNDI
LSRLDPPEAEVQIDRLITGRLQSLQTYVTQQLIRAAEIRASANLAAIKMSECVLGQSKRVDFCGKGYHLM
SFPQSAPHGVVFLHVTYVPAQEKNFTTAPAICHDGKAHFPREGVFVSNGTHWFVTQRNFYEPQIITTDNT
FVSGNCDVVIGIVNNTVYDPLQPELDSFKEELDKYFKNHTSPDVDLGDISGINASFVNIQKEIDRLNEVA
KNLNESLIDLQELGKYEQGSGYIPEAPRDGQAYVRKDGEWVLLSTFLGRSLEVLFQGPGSAWSHPQFEKG
GGSGGGGSGGSAWSHPQFEK*

**Delta (B.1.617.2) Variant 6P, 1-1208 (T19R, G142D, del156-157, R158G, L452R, T478K, D614G, P681R, D950N; 6P – F817P, A892P, A899P, A942P, K986P, V987P; Furin CS – R682G, R683S, R685S)**

MFVFLVLLPLVSSQCVNLRTRTQLPPAYTNSFTRGVYYPDKVFRSSVLHSTQDLFLPFFSNVTWFHAIHV
SGTNGTKRFDNPVLPFNDGVYFASTEKSNIIRGWIFGTTLDSKTQSLLIVNNATNVVIKVCEFQFCNDPF
LDVYYHKNNKSWMESGVYSSANNCTFEYVSQPFLMDLEGKQGNFKNLREFVFKNIDGYFKIYSKHTPINL
VRDLPQGFSALEPLVDLPIGINITRFQTLLALHRSYLTPGDSSSGWTAGAAAYYVGYLQPRTFLLKYNEN
GTITDAVDCALDPLSETKCTLKSFTVEKGIYQTSNFRVQPTESIVRFPNITNLCPFGEVFNATRFASVYA
WNRKRISNCVADYSVLYNSASFSTFKCYGVSPTKLNDLCFTNVYADSFVIRGDEVRQIAPGQTGKIADYN
YKLPDDFTGCVIAWNSNNLDSKVGGNYNYRYRLFRKSNLKPFERDISTEIYQAGSKPCNGVEGFNCYFPL
QSYGFQPTNGVGYQPYRVVVLSFELLHAPATVCGPKKSTNLVKNKCVNFNFNGLTGTGVLTESNKKFLPF
QQFGRDIADTTDAVRDPQTLEILDITPCSFGGVSVITPGTNTSNQVAVLYQGVNCTEVPVAIHADQLTPT
WRVYSTGSNVFQTRAGCLIGAEHVNNSYECDIPIGAGICASYQTQTNSRGSASSVASQSIIAYTMSLGAE
NSVAYSNNSIAIPTNFTISVTTEILPVSMTKTSVDCTMYICGDSTECSNLLLQYGSFCTQLNRALTGIAV
EQDKNTQEVFAQVKQIYKTPPIKDFGGFNFSQILPDPSKPSKRSPIEDLLFNKVTLADAGFIKQYGDCLG
DIAARDLICAQKFNGLTVLPPLLTDEMIAQYTSALLAGTITSGWTFGAGPALQIPFPMQMAYRFNGIGVT
QNVLYENQKLIANQFNSAIGKIQDSLSSTPSALGKLQNVVNQNAQALNTLVKQLSSNFGAISSVLNDILS
RLDPPEAEVQIDRLITGRLQSLQTYVTQQLIRAAEIRASANLAATKMSECVLGQSKRVDFCGKGYHLMSF
PQSAPHGVVFLHVTYVPAQEKNFTTAPAICHDGKAHFPREGVFVSNGTHWFVTQRNFYEPQIITTDNTFV
SGNCDVVIGIVNNTVYDPLQPELDSFKEELDKYFKNHTSPDVDLGDISGINASVVNIQKEIDRLNEVAKN

LNESLIDLQELGKYEQGSGYIPEAPRDGQAYVRKDGEWVLLSTFLGRSLEVLFQGPGSAWSHPQFEKGGG
SGGGGSGGSAWSHPQFEK*

**Delta (B.1.617.2) Variant 6P-Mut7, 1-1208 (T19R, G142D, del156-157, R158G, L452R, T478K, D614G, P681R, D950N; 6P – F817P, A892P, A899P, A942P, K986P, V987P; Mut7 – V705C, T883C; Furin CS – R682G, R683S, R685S)**

MFVFLVLLPLVSSQCVNLRTRTQLPPAYTNSFTRGVYYPDKVFRSSVLHSTQDLFLPFFSNVTWFHAIHV
SGTNGTKRFDNPVLPFNDGVYFASTEKSNIIRGWIFGTTLDSKTQSLLIVNNATNVVIKVCEFQFCNDPF
LDVYYHKNNKSWMESGVYSSANNCTFEYVSQPFLMDLEGKQGNFKNLREFVFKNIDGYFKIYSKHTPINL
VRDLPQGFSALEPLVDLPIGINITRFQTLLALHRSYLTPGDSSSGWTAGAAAYYVGYLQPRTFLLKYNEN
GTITDAVDCALDPLSETKCTLKSFTVEKGIYQTSNFRVQPTESIVRFPNITNLCPFGEVFNATRFASVYA
WNRKRISNCVADYSVLYNSASFSTFKCYGVSPTKLNDLCFTNVYADSFVIRGDEVRQIAPGQTGKIADYN
YKLPDDFTGCVIAWNSNNLDSKVGGNYNYRYRLFRKSNLKPFERDISTEIYQAGSKPCNGVEGFNCYFPL
QSYGFQPTNGVGYQPYRVVVLSFELLHAPATVCGPKKSTNLVKNKCVNFNFNGLTGTGVLTESNKKFLPF
QQFGRDIADTTDAVRDPQTLEILDITPCSFGGVSVITPGTNTSNQVAVLYQGVNCTEVPVAIHADQLTPT
WRVYSTGSNVFQTRAGCLIGAEHVNNSYECDIPIGAGICASYQTQTNSRGSASSVASQSIIAYTMSLGAE
NSCAYSNNSIAIPTNFTISVTTEILPVSMTKTSVDCTMYICGDSTECSNLLLQYGSFCTQLNRALTGIAV
EQDKNTQEVFAQVKQIYKTPPIKDFGGFNFSQILPDPSKPSKRSPIEDLLFNKVTLADAGFIKQYGDCLG
DIAARDLICAQKFNGLTVLPPLLTDEMIAQYTSALLAGTICSGWTFGAGPALQIPFPMQMAYRFNGIGVT
QNVLYENQKLIANQFNSAIGKIQDSLSSTPSALGKLQNVVNQNAQALNTLVKQLSSNFGAISSVLNDILS
RLDPPEAEVQIDRLITGRLQSLQTYVTQQLIRAAEIRASANLAATKMSECVLGQSKRVDFCGKGYHLMSF
PQSAPHGVVFLHVTYVPAQEKNFTTAPAICHDGKAHFPREGVFVSNGTHWFVTQRNFYEPQIITTDNTFV
SGNCDVVIGIVNNTVYDPLQPELDSFKEELDKYFKNHTSPDVDLGDISGINASVVNIQKEIDRLNEVAKN
LNESLIDLQELGKYEQGSGYIPEAPRDGQAYVRKDGEWVLLSTFLGRSLEVLFQGPGSAWSHPQFEKGGG
SGGGGSGGSAWSHPQFEK*

**Omicron (B.1.1.529) Variant 6P, 1-1208 (A67V, del69-70, T95I, G142D, del143-145, del211, ~~L212I~~, ins214EPE, G339D, S371L, S373P, S375F, K417N, N440K, G446S, S477N, T478K, E484A, Q493R, G496S, Q498R, N501Y, Y505H, T547K, D614G, H655Y, N679K, P681H, N764K, D796Y, N856K, Q954H, N969K, L981F; 6P – F817P, A892P, A899P, A942P, K986P, V987P; Furin CS – R682G, R683S, R685S)**

MFVFLVLLPLVSSQCVNLTTRTQLPPAYTNSFTRGVYYPDKVFRSSVLHSTQDLFLPFFSNVTWFHVISG
TNGTKRFDNPVLPFNDGVYFASIEKSNIIRGWIFGTTLDSKTQSLLIVNNATNVVIKVCEFQFCNDPFLD
HKNNKSWMESEFRVYSSANNCTFEYVSQPFLMDLEGKQGNFKNLREFVFKNIDGYFKIYSKHTPILVREP
EDLPQGFSALEPLVDLPIGINITRFQTLLALHRSYLTPGDSSSGWTAGAAAYYVGYLQPRTFLLKYNENG
TITDAVDCALDPLSETKCTLKSFTVEKGIYQTSNFRVQPTESIVRFPNITNLCPFDEVFNATRFASVYAW
NRKRISNCVADYSVLYNLAPFFTFKCYGVSPTKLNDLCFTNVYADSFVIRGDEVRQIAPGQTGNIADYNY
KLPDDFTGCVIAWNSNKLDSKVSGNYNYLYRLFRKSNLKPFERDISTEIYQAGNKPCNGVAGFNCYFPLR
SYSFRPTYGVGHQPYRVVVLSFELLHAPATVCGPKKSTNLVKNKCVNFNFNGLKGTGVLTESNKKFLPFQ
QFGRDIADTTDAVRDPQTLEILDITPCSFGGVSVITPGTNTSNQVAVLYQGVNCTEVPVAIHADQLTPTW
RVYSTGSNVFQTRAGCLIGAEYVNNSYECDIPIGAGICASYQTQTKSHGSASSVASQSIIAYTMSLGAEN
SVAYSNNSIAIPTNFTISVTTEILPVSMTKTSVDCTMYICGDSTECSNLLLQYGSFCTQLKRALTGIAVE
QDKNTQEVFAQVKQIYKTPPIKYFGGFNFSQILPDPSKPSKRSPIEDLLFNKVTLADAGFIKQYGDCLGD
IAARDLICAQKFKGLTVLPPLLTDEMIAQYTSALLAGTITSGWTFGAGPALQIPFPMQMAYRFNGIGVTQ
NVLYENQKLIANQFNSAIGKIQDSLSSTPSALGKLQDVVNHNAQALNTLVKQLSSKFGAISSVLNDIFSR
LDPPEAEVQIDRLITGRLQSLQTYVTQQLIRAAEIRASANLAATKMSECVLGQSKRVDFCGKGYHLMSFP
QSAPHGVVFLHVTYVPAQEKNFTTAPAICHDGKAHFPREGVFVSNGTHWFVTQRNFYEPQIITTDNTFVS
GNCDVVIGIVNNTVYDPLQPELDSFKEELDKYFKNHTSPDVDLGDISGINASVVNIQKEIDRLNEVAKNL
NESLIDLQELGKYEQGSGYIPEAPRDGQAYVRKDGEWVLLSTFLGRSLEVLFQGPGSAWSHPQFEKGGGS
GGGGSGGSAWSHPQFEK*

**Omicron (B.1.1.529) Variant 6P-Mut7, 1-1208 (A67V, del69-70, T95I, G142D, del143-145, del211, ~~L212I~~, ins214EPE, G339D, S371L, S373P, S375F, K417N, N440K, G446S, S477N, T478K, E484A, Q493R, G496S, Q498R, N501Y, Y505H, T547K, D614G, H655Y, N679K, P681H, N764K, D796Y, N856K, Q954H, N969K, L981F; 6P – F817P, A892P, A899P, A942P, K986P, V987P; Mut7 – V705C, T883C; Furin CS – R682G, R683S, R685S)**

MFVFLVLLPLVSSQCVNLTTRTQLPPAYTNSFTRGVYYPDKVFRSSVLHSTQDLFLPFFSNVTWFHVISG
TNGTKRFDNPVLPFNDGVYFASIEKSNIIRGWIFGTTLDSKTQSLLIVNNATNVVIKVCEFQFCNDPFLD
HKNNKSWMESEFRVYSSANNCTFEYVSQPFLMDLEGKQGNFKNLREFVFKNIDGYFKIYSKHTPILVREP
EDLPQGFSALEPLVDLPIGINITRFQTLLALHRSYLTPGDSSSGWTAGAAAYYVGYLQPRTFLLKYNENG
TITDAVDCALDPLSETKCTLKSFTVEKGIYQTSNFRVQPTESIVRFPNITNLCPFDEVFNATRFASVYAW
NRKRISNCVADYSVLYNLAPFFTFKCYGVSPTKLNDLCFTNVYADSFVIRGDEVRQIAPGQTGNIADYNY
KLPDDFTGCVIAWNSNKLDSKVSGNYNYLYRLFRKSNLKPFERDISTEIYQAGNKPCNGVAGFNCYFPLR
SYSFRPTYGVGHQPYRVVVLSFELLHAPATVCGPKKSTNLVKNKCVNFNFNGLKGTGVLTESNKKFLPFQ
QFGRDIADTTDAVRDPQTLEILDITPCSFGGVSVITPGTNTSNQVAVLYQGVNCTEVPVAIHADQLTPTW
RVYSTGSNVFQTRAGCLIGAEYVNNSYECDIPIGAGICASYQTQTKSHGSASSVASQSIIAYTMSLGAEN
SCAYSNNSIAIPTNFTISVTTEILPVSMTKTSVDCTMYICGDSTECSNLLLQYGSFCTQLKRALTGIAVE
QDKNTQEVFAQVKQIYKTPPIKYFGGFNFSQILPDPSKPSKRSPIEDLLFNKVTLADAGFIKQYGDCLGD
IAARDLICAQKFKGLTVLPPLLTDEMIAQYTSALLAGTICSGWTFGAGPALQIPFPMQMAYRFNGIGVTQ
NVLYENQKLIANQFNSAIGKIQDSLSSTPSALGKLQDVVNHNAQALNTLVKQLSSKFGAISSVLNDIFSR
LDPPEAEVQIDRLITGRLQSLQTYVTQQLIRAAEIRASANLAATKMSECVLGQSKRVDFCGKGYHLMSFP
QSAPHGVVFLHVTYVPAQEKNFTTAPAICHDGKAHFPREGVFVSNGTHWFVTQRNFYEPQIITTDNTFVS
GNCDVVIGIVNNTVYDPLQPELDSFKEELDKYFKNHTSPDVDLGDISGINASVVNIQKEIDRLNEVAKNL
NESLIDLQELGKYEQGSGYIPEAPRDGQAYVRKDGEWVLLSTFLGRSLEVLFQGPGSAWSHPQFEKGGGS
GGGGSGGSAWSHPQFEK*

**Mu (B.1.621) Variant 6P-Mut7, 1-1208 (T95I, Y144T, Y145S, ins146N, R346K, E484K, N501Y, D614G, P681H, D950N; 6P – F817P, A892P, A899P, A942P, K986P, V987P; Mut7 – V705C, T883C; Furin CS – R682G, R683S, R685S)**

MFVFLVLLPLVSSQCVNLTTRTQLPPAYTNSFTRGVYYPDKVFRSSVLHSTQDLFLPFFSNVTWFHAIHV
SGTNGTKRFDNPVLPFNDGVYFASIEKSNIIRGWIFGTTLDSKTQSLLIVNNATNVVIKVCEFQFCNDPF
LGVTSNHKNNKSWMESEFRVYSSANNCTFEYVSQPFLMDLEGKQGNFKNLREFVFKNIDGYFKIYSKHTP
INLVRDLPQGFSALEPLVDLPIGINITRFQTLLALHRSYLTPGDSSSGWTAGAAAYYVGYLQPRTFLLKY
NENGTITDAVDCALDPLSETKCTLKSFTVEKGIYQTSNFRVQPTESIVRFPNITNLCPFGEVFNATKFAS
VYAWNRKRISNCVADYSVLYNSASFSTFKCYGVSPTKLNDLCFTNVYADSFVIRGDEVRQIAPGQTGKIA
DYNYKLPDDFTGCVIAWNSNNLDSKVGGNYNYLYRLFRKSNLKPFERDISTEIYQAGSTPCNGVKGFNCY
FPLQSYGFQPTYGVGYQPYRVVVLSFELLHAPATVCGPKKSTNLVKNKCVNFNFNGLTGTGVLTESNKKF
LPFQQFGRDIADTTDAVRDPQTLEILDITPCSFGGVSVITPGTNTSNQVAVLYQGVNCTEVPVAIHADQL
TPTWRVYSTGSNVFQTRAGCLIGAEHVNNSYECDIPIGAGICASYQTQTNSHGSASSVASQSIIAYTMSL
GAENSCAYSNNSIAIPTNFTISVTTEILPVSMTKTSVDCTMYICGDSTECSNLLLQYGSFCTQLNRALTG
IAVEQDKNTQEVFAQVKQIYKTPPIKDFGGFNFSQILPDPSKPSKRSPIEDLLFNKVTLADAGFIKQYGD
CLGDIAARDLICAQKFNGLTVLPPLLTDEMIAQYTSALLAGTICSGWTFGAGPALQIPFPMQMAYRFNGI
GVTQNVLYENQKLIANQFNSAIGKIQDSLSSTPSALGKLQNVVNQNAQALNTLVKQLSSNFGAISSVLND
ILSRLDPPEAEVQIDRLITGRLQSLQTYVTQQLIRAAEIRASANLAATKMSECVLGQSKRVDFCGKGYHL
MSFPQSAPHGVVFLHVTYVPAQEKNFTTAPAICHDGKAHFPREGVFVSNGTHWFVTQRNFYEPQIITTDN
TFVSGNCDVVIGIVNNTVYDPLQPELDSFKEELDKYFKNHTSPDVDLGDISGINASVVNIQKEIDRLNEV
AKNLNESLIDLQELGKYEQGSGYIPEAPRDGQAYVRKDGEWVLLSTFLGRSLEVLFQGPGSAWSHPQFEK
GGGSGGGGSGGSAWSHPQFEK*

**Lambda (C.37) Variant 6P Mut7, 1-1208 (G75V, T76I, del246-252, L452Q, F490S, D614G, T859N; 6P – F817P, A892P, A899P, A942P, K986P, V987P; Mut7 – V705C, T883C; Furin CS – R682G, R683S, R685S)**

MFVFLVLLPLVSSQCVNLTTRTQLPPAYTNSFTRGVYYPDKVFRSSVLHSTQDLFLPFFSNVTWFHAIHV
SGTNVIKRFDNPVLPFNDGVYFASTEKSNIIRGWIFGTTLDSKTQSLLIVNNATNVVIKVCEFQFCNDPF
LGVYYHKNNKSWMESEFRVYSSANNCTFEYVSQPFLMDLEGKQGNFKNLREFVFKNIDGYFKIYSKHTPI
NLVRDLPQGFSALEPLVDLPIGINITRFQTLLALHDSSSGWTAGAAAYYVGYLQPRTFLLKYNENGTITD
AVDCALDPLSETKCTLKSFTVEKGIYQTSNFRVQPTESIVRFPNITNLCPFGEVFNATRFASVYAWNRKR
ISNCVADYSVLYNSASFSTFKCYGVSPTKLNDLCFTNVYADSFVIRGDEVRQIAPGQTGKIADYNYKLPD
DFTGCVIAWNSNNLDSKVGGNYNYQYRLFRKSNLKPFERDISTEIYQAGSTPCNGVEGFNCYSPLQSYGF
QPTNGVGYQPYRVVVLSFELLHAPATVCGPKKSTNLVKNKCVNFNFNGLTGTGVLTESNKKFLPFQQFGR
DIADTTDAVRDPQTLEILDITPCSFGGVSVITPGTNTSNQVAVLYQGVNCTEVPVAIHADQLTPTWRVYS
TGSNVFQTRAGCLIGAEHVNNSYECDIPIGAGICASYQTQTNSPGSASSVASQSIIAYTMSLGAENSCAY
SNNSIAIPTNFTISVTTEILPVSMTKTSVDCTMYICGDSTECSNLLLQYGSFCTQLNRALTGIAVEQDKN
TQEVFAQVKQIYKTPPIKDFGGFNFSQILPDPSKPSKRSPIEDLLFNKVTLADAGFIKQYGDCLGDIAAR
DLICAQKFNGLNVLPPLLTDEMIAQYTSALLAGTICSGWTFGAGPALQIPFPMQMAYRFNGIGVTQNVLY
ENQKLIANQFNSAIGKIQDSLSSTPSALGKLQDVVNQNAQALNTLVKQLSSNFGAISSVLNDILSRLDPP
EAEVQIDRLITGRLQSLQTYVTQQLIRAAEIRASANLAATKMSECVLGQSKRVDFCGKGYHLMSFPQSAP
HGVVFLHVTYVPAQEKNFTTAPAICHDGKAHFPREGVFVSNGTHWFVTQRNFYEPQIITTDNTFVSGNCD
VVIGIVNNTVYDPLQPELDSFKEELDKYFKNHTSPDVDLGDISGINASVVNIQKEIDRLNEVAKNLNESL
IDLQELGKYEQGSGYIPEAPRDGQAYVRKDGEWVLLSTFLGRSLEVLFQGPGSAWSHPQFEKGGGSGGGG
SGGSAWSHPQFEK*

**Reference Sequence (Wuhan-Hu-1, 1-1273); UniProt# P0DCT2**

>QHD43416.1 surface glycoprotein [severe acute respiratory syndrome coronavirus 2]
MFVFLVLLPLVSSQCVNLTTRTQLPPAYTNSFTRGVYYPDKVFRSSVLHSTQDLFLPFFSNVTWFHAIHV
SGTNGTKRFDNPVLPFNDGVYFASTEKSNIIRGWIFGTTLDSKTQSLLIVNNATNVVIKVCEFQFCNDPF
LGVYYHKNNKSWMESEFRVYSSANNCTFEYVSQPFLMDLEGKQGNFKNLREFVFKNIDGYFKIYSKHTPI
NLVRDLPQGFSALEPLVDLPIGINITRFQTLLALHRSYLTPGDSSSGWTAGAAAYYVGYLQPRTFLLKYN
ENGTITDAVDCALDPLSETKCTLKSFTVEKGIYQTSNFRVQPTESIVRFPNITNLCPFGEVFNATRFASV
YAWNRKRISNCVADYSVLYNSASFSTFKCYGVSPTKLNDLCFTNVYADSFVIRGDEVRQIAPGQTGKIAD
YNYKLPDDFTGCVIAWNSNNLDSKVGGNYNYLYRLFRKSNLKPFERDISTEIYQAGSTPCNGVEGFNCYF
PLQSYGFQPTNGVGYQPYRVVVLSFELLHAPATVCGPKKSTNLVKNKCVNFNFNGLTGTGVLTESNKKFL
PFQQFGRDIADTTDAVRDPQTLEILDITPCSFGGVSVITPGTNTSNQVAVLYQDVNCTEVPVAIHADQLT
PTWRVYSTGSNVFQTRAGCLIGAEHVNNSYECDIPIGAGICASYQTQTNSPRRARSVASQSIIAYTMSLG
AENSVAYSNNSIAIPTNFTISVTTEILPVSMTKTSVDCTMYICGDSTECSNLLLQYGSFCTQLNRALTGI
AVEQDKNTQEVFAQVKQIYKTPPIKDFGGFNFSQILPDPSKPSKRSFIEDLLFNKVTLADAGFIKQYGDC
LGDIAARDLICAQKFNGLTVLPPLLTDEMIAQYTSALLAGTITSGWTFGAGAALQIPFAMQMAYRFNGIG
VTQNVLYENQKLIANQFNSAIGKIQDSLSSTASALGKLQDVVNQNAQALNTLVKQLSSNFGAISSVLNDI
LSRLDKVEAEVQIDRLITGRLQSLQTYVTQQLIRAAEIRASANLAATKMSECVLGQSKRVDFCGKGYHLM
SFPQSAPHGVVFLHVTYVPAQEKNFTTAPAICHDGKAHFPREGVFVSNGTHWFVTQRNFYEPQIITTDNT
FVSGNCDVVIGIVNNTVYDPLQPELDSFKEELDKYFKNHTSPDVDLGDISGINASVVNIQKEIDRLNEVA
KNLNESLIDLQELGKYEQYIKWPWYIWLGFIAGLIAIVMVTIMLCCMTSCCSCLKGCCSCGSCCKFDEDD
SEPVLKGVKLHYT*

**Key:**

Foldon, T4 fibritin trimerization domain [S10]
HRV3C protease cleavage sequence (modified from [S11])
TwinStrep tag for affinity purification [S12]
8x His-tag
Possible conflict
* stop codon
